# Serotype-independent inhibition of *S. pneumoniae* by SMiTE, a commensal-derived bacteriocin

**DOI:** 10.64898/2026.02.25.707991

**Authors:** João Lança, Joana Bryton, João Borralho, Catarina Candeias, Wilson Antunes, Amir Pandi, Raquel Sá-Leão

## Abstract

*Streptococcus pneumoniae* remains a leading cause of disease despite widespread vaccination, highlighting the need for serotype-independent strategies. We recently identified commensal streptococci that produce bacteriocins with anti-pneumococcal activity. Here, we evaluate these bacteriocins as candidates for pneumococcal control. Using cell-free protein synthesis, we screened 58 bacteriocins, the majority of which absent in available pneumococcal genomes, and identified SMiTE as the most potent. Purified SMiTE disrupted pneumococcal membrane integrity, as shown by confocal, transmission, and scanning electron microscopy, induced ATP leakage, and triggered transcriptional responses consistent with envelope stress and metabolic remodeling. In a mouse nasopharyngeal colonization model, intranasal SMiTE treatments reduced pneumococcal loads by 65-fold with no significant weight loss. SMiTE inhibited a broad range of serotypes, with strongest activity against serotype 3, which is poorly controlled by current vaccines, while sparing most oral and upper respiratory tract commensals. Repeated sub-inhibitory exposure did not select resistant mutants in line with the membrane-targeting mechanism. These findings establish SMiTE as a commensal-derived strategy for serotype-independent, microbiota-sparing pneumococcal decolonization.

## Introduction

*Streptococcus pneumoniae*, or the pneumococcus, remains a major global health threat. This opportunistic pathogen is responsible for a wide range of infections including otitis media, pneumonia, bacteremia, and meningitis^1,2^. In 2019, it was the third leading cause of deaths associated with infectious diseases worldwide and was responsible for the highest number of years of life lost from bacterial infections^3,4^. Asymptomatic colonization of the upper respiratory tract is a prerequisite for disease and a reservoir for transmission. Therefore, targeting colonization is a promising strategy to reduce pneumococcal burden^5,6^.

Current strategies to prevent and control pneumococcal disease, including antibiotics and pneumococcal conjugate vaccines (PCVs), while effective, face limitations. Antibiotics, while essential for treatment, are often associated with disruption of the healthy microbiota and select for antimicrobial resistance^7,8^. PCVs, despite highly effective at preventing disease, have limited valency targeting up to 21 serotypes of the more than 100 described so far, and often lead to serotype replacement in both colonization and disease^9–11^. Thus, new interventions that selectively target pneumococcal colonization across serotypes and complement the existing ones are needed. In response to these limitations, a diverse range of alternative approaches is being explored. These strategies aim to broaden vaccine coverage, modulate host-pathogen interactions, or directly target pneumococcal survival and colonization using non-traditional antimicrobial therapies^12,13^. Among the latter, bacteriocins represent a promising but underexplored class of targeted antimicrobials^14^.

Bacteriocins are ribosomally synthesized antimicrobial peptides produced by bacteria that frequently inhibit closely related species, often by disrupting membranes, leading to rapid cell death^15,16^. These antimicrobial peptides are also considered to be less prone to resistance development^17,18^. Some bacteriocins, such as nisin and pediocin PA-1, are already used in food preservation or are under preclinical development for human infections, respectively^19–23^. Although no bacteriocin has yet been approved for therapeutic use, multiple candidates are actively being explored. For example, R-pyocins have been shown to prevent and treat acute pneumonia caused by a high-risk clone of *Pseudomonas aeruginosa* in a murine model^24^. To our knowledge, bacteriocins alone have not been explored as a strategy to selectively reduce *S. pneumoniae* colonization.

Recently, we identified seven commensal streptococci (*S. oralis* strain A22 and *S. mitis* strains B22 to G22) with serotype-independent inhibitory activity against *S. pneumoniae*^14^. Across these strains, a total of 61 bacteriocin loci were identified, of which 35 encoded one or more bacteriocins, encompassing bacteriocin-like peptide (*blp*), competence-associated bacteriocin (*cab*), streptococcin, and lantibiotic families (**Supplementary Fig. S1**). Individual strains encoded between three and eleven bacteriocin loci, comprising nine to 23 putative bacteriocins and eleven to 26 immunity proteins. Notably, approximately 80% of the bacteriocins and 92% of the immunity proteins were absent from a collection of over 7,000 pneumococcal genomes, underscoring the distinct and largely unexplored nature of this commensal streptococcal bacteriocin repertoire. Construction of deletion mutants of 28 of the bacteriocin loci, resulted in partial or complete loss of inhibitory phenotype in 14 mutants under at least one condition tested. These findings strongly suggest that bacteriocins are major contributors to the observed anti-pneumococcal activity. However, the individual contribution of specific bacteriocins remained unresolved, as each locus typically encodes multiple co-expressed peptides^14^.

Here, we used cell-free protein synthesis (CFPS) to synthesize the 58 bacteriocins encoded by strains A22 to G22 and evaluate their anti-pneumococcal activity. This approach addresses key methodological limitations, as these peptides are often difficult to obtain through chemical synthesis or recombinant production. We show that several display inhibitory activities and one, in particular, here named SMiTE, has consistent and potent activity. We found that SMiTE disrupts membrane integrity, induces midcell and surface deformations, and causes ATP leakage. In a mouse nasopharyngeal colonization model, SMiTE significantly reduced pneumococcal burden. Importantly, SMiTE displayed serotype-independent activity across multiple epidemiologically relevant pneumococcal serotypes and limited off-target effects on other bacteria. These results position SMiTE as a promising naturally occurring antimicrobial agent for serotype-independent pneumococcal decolonization, with potential to complement existing interventions aimed at reducing disease burden.

## Results

### Cell-free protein synthesis enables functional screening of bacteriocins with anti-pneumococcal activity

In our previous work, we identified seven commensal streptococcal strains (one *S. oralis* A22 and six *S. mitis* B22 to G22), isolated from healthy individuals, with serotype-independent inhibitory activity against *S. pneumoniae*^14^. Construction of deletion mutants confirmed that bacteriocin loci were largely responsible for this phenotype. However, the specific bacteriocins mediating the inhibitory activity remained unclear, as several of the loci identified encode multiple putative bacteriocins (**Supplementary Fig. S1**). Moreover, one of the seven commensal strains was not genetically transformable (*S. mitis* E22), limiting functional interrogation of its bacteriocin loci. Genomic analysis of strains A22 to G22 identified, overall, 70 putative bacteriocin genes associated with bacteriocin-like peptide (*blp*), competence-associated (*cab*), streptococcin (*scc*) and lantibiotic (*lan*) loci. Remarkably, several of these genes and its putative cognate immunity proteins were completely absent from a collection of 7,548 pneumococcal genomes^14^.

To gain insights on the activity of the putative bacteriocins previously identified we resorted to cell-free protein synthesis. We focused on bacteriocins predicted not to require post-translational modifications, *i.e.*, those encoded in *blp*, *cab*, and *scc* loci, which allowed for direct synthesis and activity screening. Based on sequence and structure predictions, *blp*-encoded bacteriocins typically adopt a helix-turn-helix α-helical conformation and range from 35 to 61 amino acids in length. In contrast, *cab* bacteriocins were generally smaller, ranging from 20 to 53 amino acids, and predicted to form a single α-helix. Streptococcins were predicted to adopt a compact β-sandwich fold composed of antiparallel β-sheets, similar to lactococcin 972, and are larger, ranging from 64 to 75 amino acids (**Supplementary Fig. S2**).

In total, using CFPS, we synthesized 58 bacteriocins and screened their activity against *S. pneumoniae* P537 and D39 strains (**Fig. 1, Supplementary Fig. S2**). Seven bacteriocins - Bac2v1, BlpK_1_, BlpK_2_, BlpK_3_, BlpOl_ike1_, BlpO_like2_, and BlpW_1_ - significantly inhibited pneumococcal growth in planktonic culture, primarily by delaying entry into the exponential phase (**Fig. 1A-B, Supplementary Fig. S3A-B**). Despite only minor sequence variations, some bacteriocins displayed markedly different levels of activity. Notably, BlpO_like2_ differs from BlpO_like1_ by a single amino acid, while BlpK_2_ and BlpK_3_ differ from BlpK_1_ by four and three amino acids, respectively (**Supplementary Fig. S3C**). Furthermore, eight bacteriocins exhibited activity against *S. pneumoniae* P537 and/or D39 on solid agar (**Fig. 1C-D, Supplementary Fig. S3D-E**). Among all candidates, BlpO_like2_ - hereinafter named SMiTE (*<u>S</u>. <u>mi</u>tis* toxin <u>e</u>ffector) - consistently inhibited both pneumococcal strains in both assays.

**Fig. 1.**
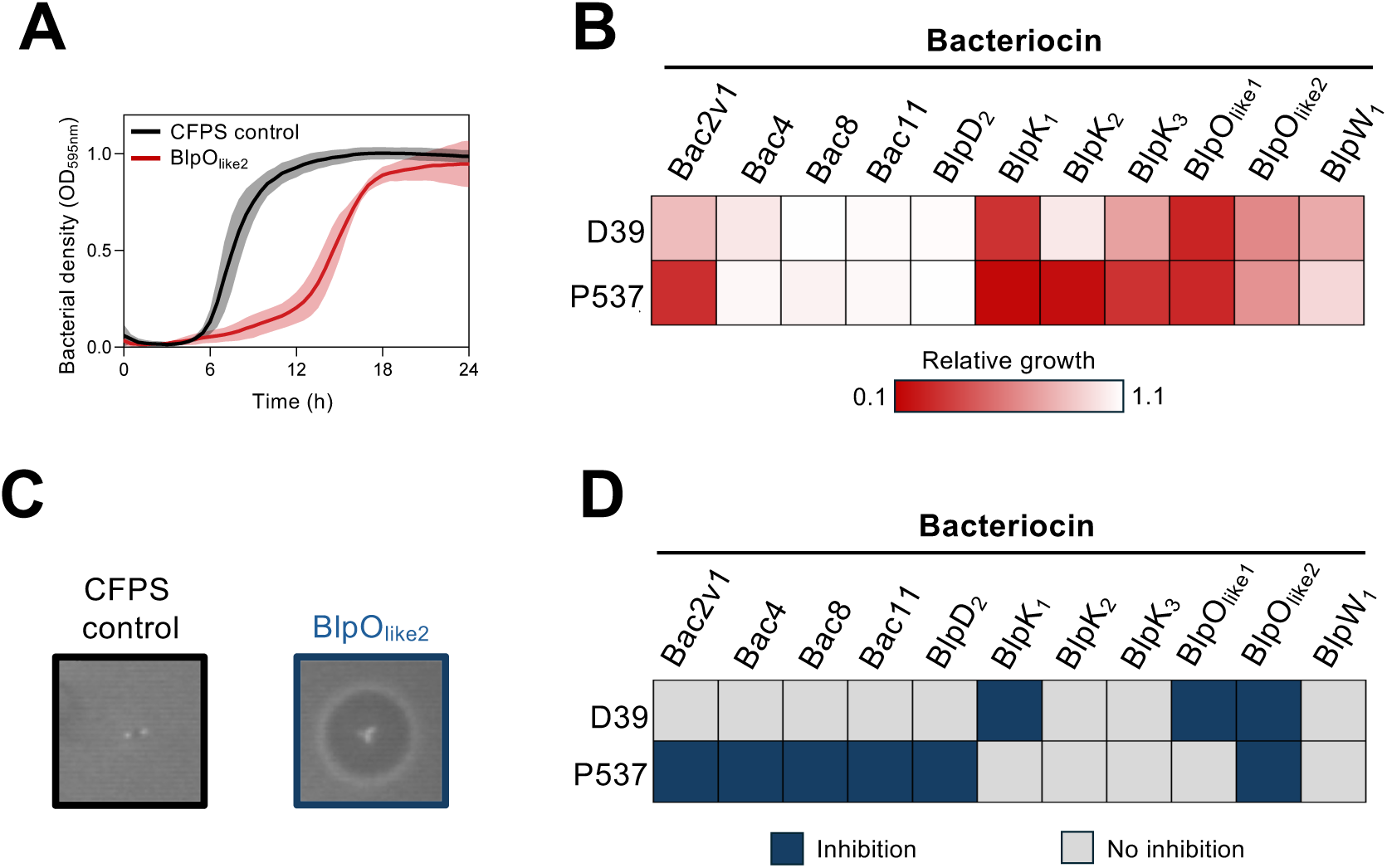
SMiTE (BlpO_like2_) has consistent inhibitory activity against *S. pneumoniae*. (A) Representative growth curve of *S. pneumoniae* D39 exposed to cell-free synthesized SMiTE (BlpO_like2_) (red) or cell-free protein synthesis (CFPS) control (black). Lines represent means and shaded areas indicate standard deviations of three independent experiments. (B) Summary of planktonic growth inhibition by bacteriocins that affected at least one *S. pneumoniae* strain in either assay. Color intensity reflects relative growth (AUC of bacteriocin-exposed culture / AUC of CFPS control culture), with redder cells indicating stronger inhibition. (C) Representative images of solid media assay showing absence of inhibition with CFPS control and an inhibition halo with SMiTE (BlpO_like2_) against *S. pneumoniae* D39. (D) Summary of inhibition halos observed on solid media for bacteriocins that inhibited at least one *S. pneumoniae* in either assay. Blue cells indicate presence of an inhibition halo, while grey cells indicate absence. All assays were performed in triplicate.

Together, these assays identified various bacteriocins with measurable inhibitory activity against *S. pneumoniae* across planktonic and solid culture assays. Among these, we selected SMiTE for further characterization.

### SMiTE disrupts *S. pneumoniae* membranes causing leakage of intracellular contents

Bacteriocins are commonly described as antimicrobial peptides that act primarily by disrupting bacterial membranes, often through pore formation that compromises membrane integrity and leads to cell death ^25^. SMiTE is a 55-amino acid class II bacteriocin ^25^ predicted to adopt a helix-turn-helix α-helical structure and contains four GxxxG and eight GxxxG-like motifs, highlighting its propensity for multimerization (**Fig. 2A-B**). Sequence-based analysis revealed a strong enrichment in non-polar aliphatic residues (65.5%), together with a moderate content of aromatic residues (10.9%), consistent with a highly hydrophobic primary structure. The peptide exhibits a very high aliphatic index (99.64), a parameter associated with enhanced thermal stability, and a strongly positive GRAVY score (0.774), reflecting a propensity for interaction with lipid bilayers. In addition, the low instability index (12.74) suggests that SMiTE is intrinsically stable under physiological conditions (**Supplementary Table S1**).

**Fig. 2.**
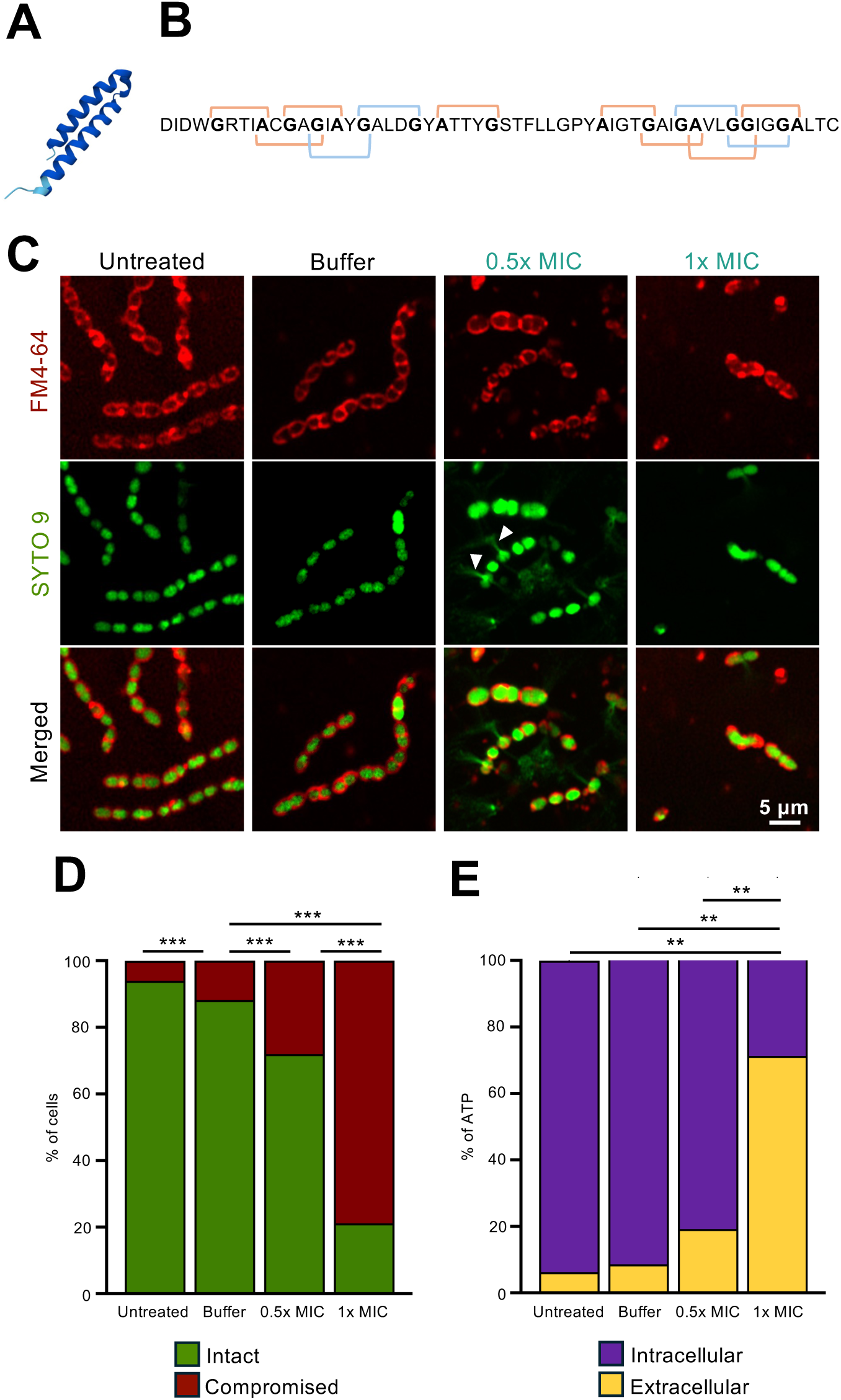
SMiTE compromises pneumococcal membrane integrity in a dose-dependent manner. (A) Predicted structure of SMiTE monomer generated by Alphafold (ColabFoldjj v1.5.5) and is colored according to model confidence using the predicted local distance test (pLDDT): light blue indicates confident predictions (70 < pLDDT < 90), and dark blue indicates very high confident predictions (pLDDT > 90). (B) Sequence map of SMiTE highlighting of GxxxG (blue) and GxxxG-like (orange) predicted to contribute to multimer formation. (C) Representative confocal fluorescence images of *S. pneumoniae* D39 cultures untreated, exposed to the bacteriocin buffer and to 0.5x MIC or 1x MIC of SMiTE. Cultures were stained with FM4-64 (membrane) and SYTO9 (nucleic acid). White arrows indicate DNA leakage. (D) LIVE/DEAD assay. Quantification of membrane permeability, showing the percentage of cells stained with propidium iodine (permeabilized, red) or SYTO9 (intact, green) under each treatment condition. A total of 6,247 cells (THY), 5,715 cells (buffer), 6,892 cells (0.5x MIC), and 579 cells (1x MIC) were analyzed; the lower cell number at 1x MIC reflects the overall reduced cell abundance under this condition. A binomial GML with Tukey’s HSD post-hoc test was used to compare the proportion of live cells. *** *P* < 0.001. (E) Relative intracellular and extracellular ATP levels in untreated, treated with bacteriocin buffer, 0.5x MIC or 1x MIC of SMiTE, demonstrating dose-dependent leakage of intracellular components. The Kruskal-Wallis test followed by pairwise Wilcoxon post-hoc tests was used to compare differences in extracellular ATP. ** *P* < 0.01. All experiments were performed in triplicate.

To investigate its mechanism of action, SMiTE was heterologously expressed and purified using immobilized metal affinity chromatography (IMAC). Western blot analysis using an anti-His-tag antibody revealed multiple bands in the purified fraction, with molecular weights corresponding to approximately 13, 26, and 39 kDa, which we hypothesize correspond to dimeric, tetrameric and hexameric forms of SMiTE, respectively (**Supplementary Fig. S4A**). Based on these observations, structural models of the SMiTE multimers were generated using AlphaFold (**Supplementary Fig. S4B**). The predicted multimers exhibit electrostatic surface patches that may facilitate membrane insertion and pore-forming mechanism (**Supplementary Fig. S4C**). Finally, the minimum inhibitory concentration (MIC) of purified SMiTE against *S. pneumoniae* D39, determined by microdilution method following EUCAST guidelines, was 108 nM.

We next examined the effect of SMiTE on pneumococcal membrane integrity. First, D39 planktonic cultures were treated with 0.5x MIC or 1x MIC of SMiTE or buffer alone, then stained with FM4-64 to label the membrane and SYTO9 to label nucleic acids prior to confocal fluorescence microscopy. Untreated and buffer-treated cells formed long, well-organized diplococcal chains with smooth and uniform membranes. In contrast, SMiTE-treated cells displayed morphological abnormalities, including distortion and irregularities at the division septum, indicative of cell envelope disruption. Notably, SYTO9 fluorescence was detected outside 0.5x MIC-treated cells, indicating leakage of intracellular DNA and compromised membrane integrity. (**Fig. 2C**).

To further assess membrane permeabilization, we performed LIVE/DEAD staining coupled with confocal microscopy, and monitored SYTO9 uptake, which permeates all cells, and propidium iodide, which selectively enters and stains cells with compromised membranes.

LIVE/DEAD imaging revealed a clear dose-dependent increase in membrane permeabilization following SMiTE treatment, indicating a progressive loss of membrane integrity and cellular viability. Approximately 30% and 80% of cells exhibited compromised membranes at 0.5x MIC and 1x MIC, respectively, compared to ∼6% and ∼12% in untreated and buffer-treated controls, respectively (**Fig. 2D and Fig. S5**).

Finally, to complement this assay, we quantified intracellular and extracellular ATP levels as an additional indicator of membrane leakage and cell viability. Consistent with the previous results, ATP quantification showed a parallel dose-response: extracellular ATP rose to approximately 20% and 70% at 0.5x MIC and 1x MIC, respectively, whereas untreated and buffer-treated cultures showed only ∼6% and ∼9% extracellular ATP (**Fig. 2E**).

Together, these findings indicate that SMiTE compromises pneumococcal membrane integrity in a dose-dependent manner, leading to increased permeability and leakage of intracellular contents such as ATP, consistent with a substantial fraction of the population undergoing lethal damage at higher SMiTE concentrations. These results establish membrane disruption as a central mechanism underlying the anti-pneumococcal activity of SMiTE, consistent with the mode of action described for other clinically relevant bacteriocins.

### SMiTE causes septal damage and pore-like disruptions in *S. pneumoniae*

Because many membrane-targeting bacteriocins kill by forming defined pores or inducing localized disruptions of the cell envelope, we employed transmission electron microscopy (TEM) and scanning electron microscopy (SEM) to investigate whether SMiTE produces structural alterations in *S. pneumoniae*.

Low-magnification TEM micrographs of SMiTE-treated cultures revealed an accumulation of cellular debris, including fragmented membranes and dispersed capsular material (**Fig. 3A**). In contrast, cultures treated with the bacteriocin buffer alone showed little to no cellular debris, indicating that these effects were specific to SMiTE exposure. At higher magnification, deformations were frequently observed at the midcell region. Quantitative analysis of septal damage showed that 24.9% of cells at 0.5x MIC and 26.9% at 1x MIC exhibited septum defects, compared to only 8% in buffer-treated cells, indicating that SMiTE preferentially disrupts the division site (**Fig. 3B**).

**Fig. 3.**
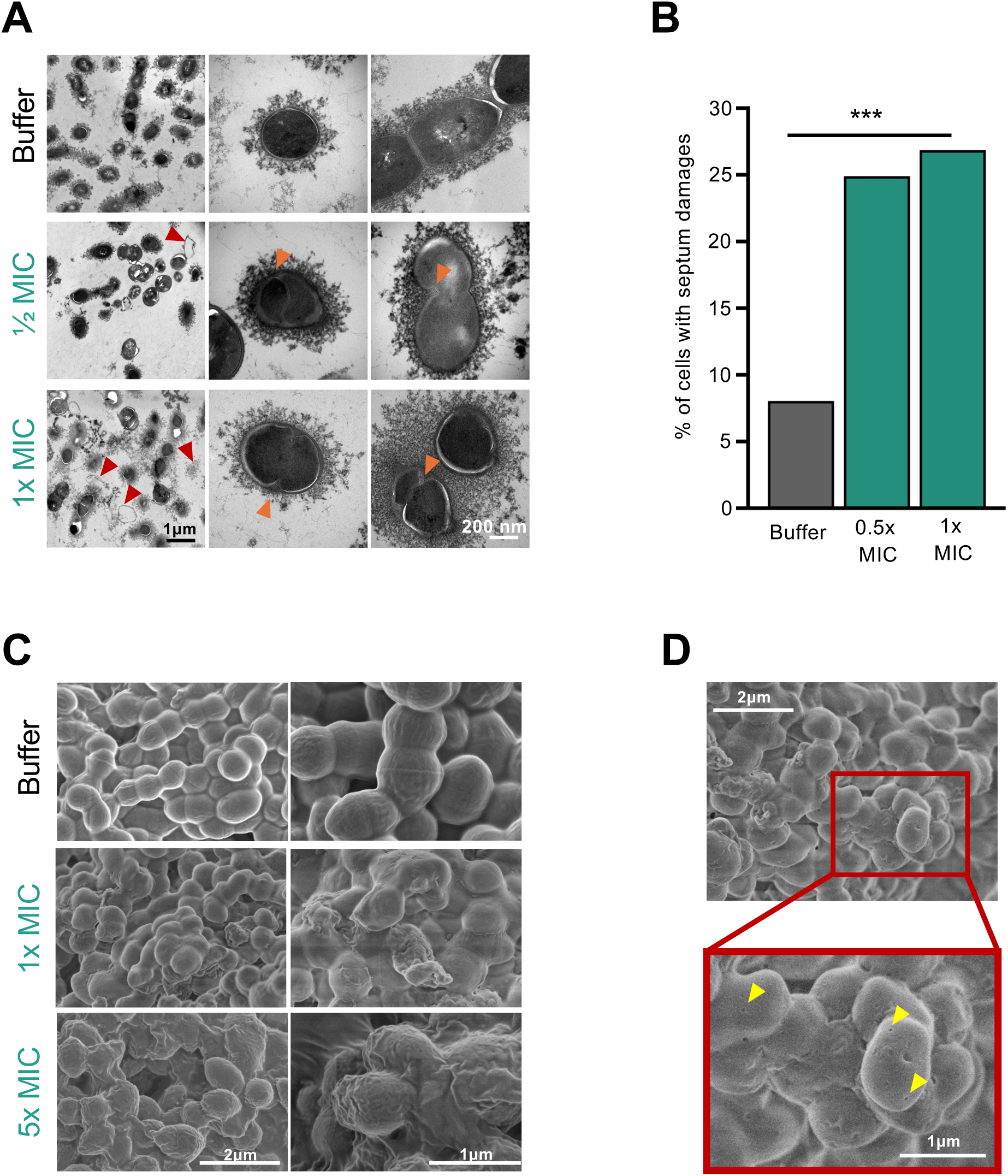
SMiTE induces structural damages in *S. pneumoniae*. (A) Representative transmission electron microscopy (TEM) images of *S. pneumoniae* D39 exposed to the bacteriocin buffer, ½ MIC, or MIC of SMiTE. Cellular debris is indicated by red arrows and deformations at the midcell region are indicated by orange arrows. (B) Blinded quantification of cells with septal damages on TEM images. Images were analyzed independently by two observers, who quantified the total number of cells and the number of cells exhibiting septal damage; because counts were performed independently, slight differences in total cell numbers were observed. The numbers of cells analyzed by observer 1 and observer 2, respectively, were: buffer, 137 and 144; 0.5x MIC, 158 and 162; and 1x MIC, 190 and 158. A binomial generalized linear mixed-effects model (GLMM) with *Condition* as a fixed effect and *Observer* as a random effect was used to compare the proportion of cells with septal damages between conditions. *** *P* < 0.001. (C) Representative scanning electron microscopy (SEM) images of *S. pneumoniae* D39 exposed to the bacteriocin buffer, 0.5x MIC, or 1x MIC of SMiTE, showing dose-dependent surface alterations. (D) High-magnification SEM of cells treated with MIC of SMiTE, with pore-like structures indicated by yellow triangles. TEM and SEM experiments were performed once.

In SEM, cells treated with the bacteriocin buffer alone maintained smooth, spherical-to-oval shapes and formed orderly and elongated diplococcal chains (**Fig. 3C**). In contrast, SMiTE treatment led to disorganized chains with closely packed cells, consistent with a stress response. At 1x MIC, many cells were surrounded by a dense matrix-like material, likely comprising leaked intracellular components. This phenotype became even more pronounced at 5x MIC, where the surface of treated cells appeared more textured and rugged, likely reflecting deformation of the cell surface (**Fig. 3C**). Higher-magnification images of 1x MIC-treated cells revealed surface discontinuities resembling pore-like structures, further supporting the hypothesis that SMiTE disrupts the pneumococcal cell membrane (**Fig. 3D**).

Together, TEM and SEM analysis demonstrate that SMiTE causes severe, concentration-dependent damage on *S. pneumoniae*. TEM revealed internal membrane deformations and septal defects, while SEM highlighted surface distortions, disorganized chains, and pore-like structures at higher magnifications. These findings collectively support a membrane-targeting mechanism of action, consistent with pore-forming bacteriocins that compromise bacterial integrity and drive cell death.

### SMiTE triggers metabolic and regulatory remodeling in *S. pneumoniae*

To investigate the transcriptional response of *S. pneumoniae* to SMiTE, RNA-Seq was performed on cultures treated with 0.5x MIC of SMiTE or bacteriocin buffer as a control. The data were analyzed through differential expression analysis, gene set enrichment analysis (GSEA), and gene ontology (GO) enrichment to capture transcriptional changes at the level of individual genes, coordinated gene sets, and broader functional categories, respectively.

Differential expression analysis revealed a marked transcriptional reprogramming upon SMiTE exposure. Fifteen genes were significantly upregulated, including four genes (*glnH*, *glnQ*, *glnPa* and *glnPb*) within a single operon involved in glutamine transport. Five genes were significantly downregulated upon SMiTE treatment: an acyl protein carrier protein (*acpP*), a cysteine synthase (*cysK*), a class I glutamine aminotransferase (*yvdE*), a hypothetical protein (SPD_1360), and an IS66 family transposase (SPD_2038) (**Figure 4A-B, Table 1 and Supplementary Table S2**).

**Fig. 4.**
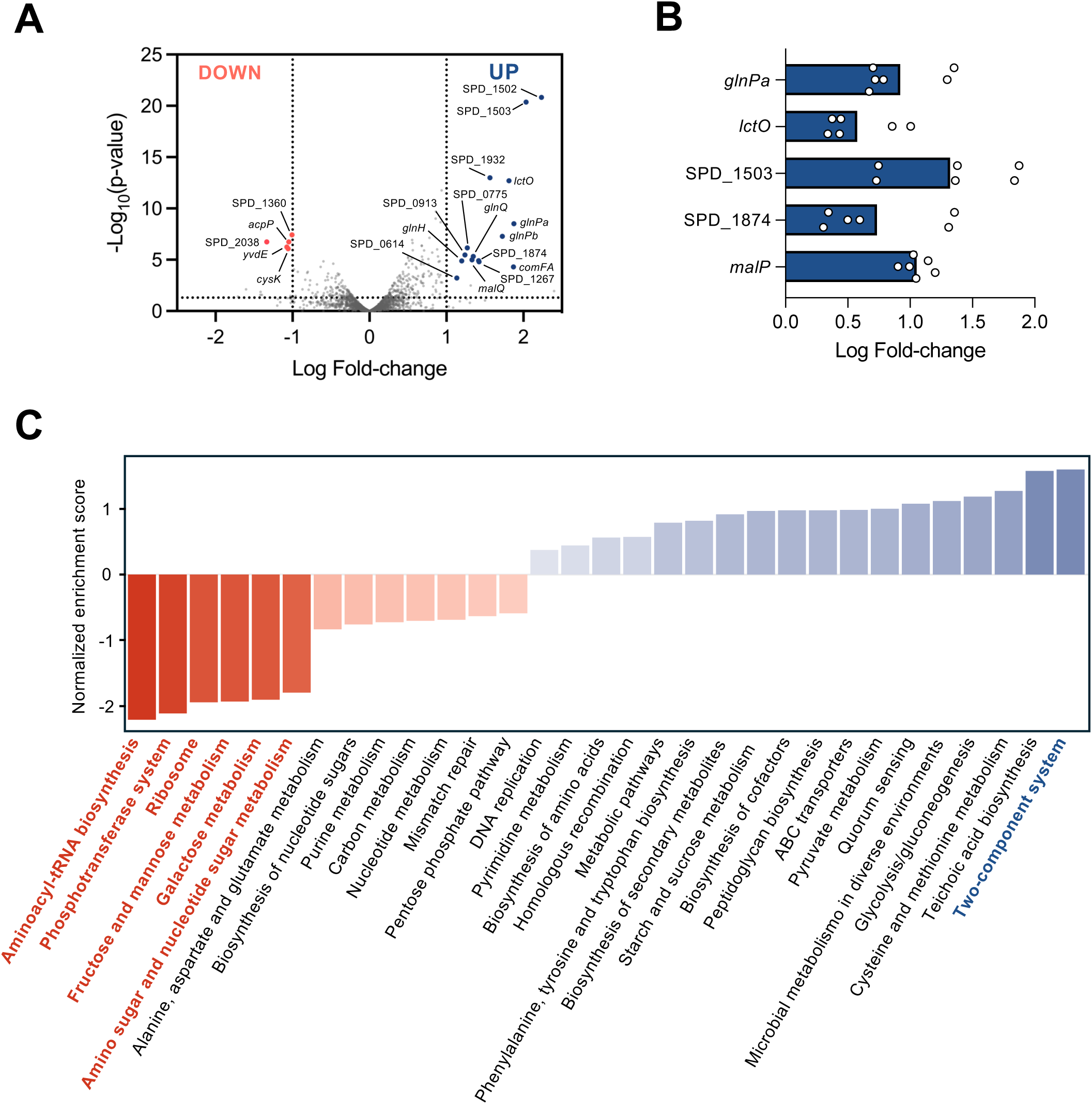
SMiTE triggers transcriptional reprogramming and metabolic adaptation in *S. pneumoniae*. (A) Volcano plot of differentially expressed genes (DEGs) in *S. pneumoniae* D39 treated with 0.5x MIC of SMiTE versus bacteriocin buffer. DEGs were considered if false discovery rate (FDR)<0.05, *P*<0.05 and log fold-change (logFC)>1 (upregulated) or logFC<1 (downregulated). Significantly downregulated genes are shown in red, upregulated are shown in blue, and non-significant genes in gray. (B) RT-qPCR validation of RNA-Seq results for five selected genes. Fold-changes were calculated using the 2^-ΔΔCt^ method and normalized to the housekeeping gene *gyrA*. (C) Summary of pathway-level changes identified by gene set enrichment analysis (GSEA). Pathways with significant upregulation or downregulation are highlighted in color.

**Table 1.** Differentially expressed genes in *S. pneumoniae* D39 exposed to SMiTE.

| Gene ID | Gene name<sup>a</sup> | Function<sup>a</sup> | MIC relative to wt |
| --- | --- | --- | --- |
| **UPREGULATED** |  |  |  |
| SPD_0614 | - | ABC transporter ATP-binding protein | 2x |
| SPD_0615 | *glnH* | Glutamine ABC transporter substrate-binding protein | = |
| SPD_0616 | *glnQ* | Glutamine ABC transporter ATP-binding protein | = |
| SPD_0617 | *glnPb* | Glutamine ABC transporter permease | = |
| SPD_0618 | *glnPa* | Glutamine ABC transporter permease | = |
| SPD_0619 | *lctO* | L-Lactate 2-monooxygenase | 2x |
| SPD_0775 | - | Acetyltransferase | = |
| SPD_0913 | - | Extracellular protein | = |
| SPD_1267 | - | Duplicated ATPase component of energizing module of putative ECF transporter | 2x |
| SPD_1502 | - | ABC transporter substrate-binding protein – sugar transport | = |
| SPD_1503 | - | Putative beta-D-galactosidase | = |
| SPD_1874 | - | Surface immunogenic protein, LysM domain-containing protein, +Q8033 (Putative) N-acetylmuramidase | 2x |
| SPD_1932 | *malP* | Maltodextrin phosphorylase | 0.5x |
| SPD_1933 | *malQ* | 4-alpha-glucanotransferase (amylomaltase) | 0.5x |
| SPD_2035 | *comFA* | DNA transporter ATPase, ComF operon protein A | = |
| **DOWNREGULATED** |  |  |  |
| SPD_0381 | *acpP* | Acyl carrier protein | Not tested |
| SPD_1360 | - | Hypothetical protein | = |
| SPD_1899 | *yvdE* | Glutamine aminotransferase, class I | 2x |
| SPD_2037 | *cysK* | Cysteine synthase | 2x |
| SPD_2038 | - | IS66 family transposase, Orf3, truncation | 2x |

<sup>a</sup>Gene name and function were obtained from PneumoBrowse 2<sup>59</sup>.

At the pathway level, GSEA identified one significantly upregulated gene set annotated as two-component regulatory systems, and six significantly downregulated pathways, including aminoacyl-tRNA biosynthesis, phosphotransferase systems, ribosomal components and multiple carbohydrate metabolism pathways (fructose/mannose, galactose, and amino sugar/nucleotide sugar metabolism) (**Fig. 4C and Supplementary Tables S3 and S4**). Within these gene sets and pathways, specific genes contributed most strongly to the enrichment signal. For example, *comB*, *vraR*, and *pstS* were among the top contributors in the two-component system; *tRNA-Arg-5*, *tRNA-Val-3* and *tRNA-Arg-4* in aminoacyl-tRNA biosynthesis; SPD_0559, SPD_0560 and *gadW* in phosphotransferase system; *rpmB*, *rpmH* and *rpsP* in ribosome; *gadW*, *fucK* and *gadE* in fructose and mannose metabolism; *bgaC*, SPD_0295 and *lacA* in galactose metabolism; and *gadW*, *gadE* and *manN* in amino sugar and nucleotide sugar metabolism (**Supplementary Fig. S4 and Supplementary Table S4**). These results indicate coordinated shifts in regulatory, translational, and metabolic processes.

GO enrichment analysis of the core genes identified by GSEA further supported these observations (**Supplementary Figure S7 and Supplementary Tables S4 and S5**). In the molecular function category, enriched GO terms were primarily associated with ribosome-related functions and sugar uptake and carbohydrate metabolism, consistent with alterations in protein synthesis and metabolic activity. Similarly, enrichment of GO terms within the biological processes category highlighted pathways related to carbohydrate metabolism and translation, indicating broad metabolic and translational reprogramming. Notably, the lipoteichoic acid biosynthesis process, which is involved in cell envelope remodeling, was also overrepresented. In the cellular component category, GO terms associated with the cytosolic small and large ribosomal subunit, as well as the plasma membrane, were enriched, supporting the hypothesis that SMiTE perturbs the cell envelope and triggers membrane-related stress responses.

To assess the functional contribution of differentially expressed genes (DEGs) to SMiTE susceptibility, deletion mutants were constructed for all DEGs. For genes located within the same operon, the entire operon was deleted. A total of 15 mutants were constructed. The *acpP* deletion mutant did not grow in the medium used for all assays and was therefore excluded from further analysis. All remaining deletion mutants reached stationary phase within 8 h of inoculation, although some exhibited minor variations in growth. These observations indicate that the differences detected in MIC values are unlikely to be due to major discrepancies in growth (**Supplementary Fig. S8**). MIC evaluation revealed that deletion of SPD_0614, *lctO*, SPD_1874, *yvdE*, and *cysK* increased the MIC twofold, whereas deletion of *malPQ* reduced the MIC by half (**Table 1**). These results demonstrate that several DEGs contribute directly to SMiTE susceptibility and highlight specific metabolic and transport pathways that modulate bacteriocin sensitivity.

Collectively, these transcriptomic analyses demonstrate that *S. pneumoniae* mounts a multifaceted response to sub-inhibitory concentrations of SMiTE. Transcriptomic changes suggest metabolic reprogramming and regulatory adaptation, while functional validation with deletion mutants confirmed that several differentially expressed genes directly influence SMiTE susceptibility. Enrichment of membrane-associated processes aligns with microscopy observations of cell envelope perturbation, supporting the conclusion that SMiTE exposure triggers a coordinated stress response in *S. pneumoniae* integrating both metabolic and regulatory remodeling.

### SMiTE decreases *S. pneumoniae* colonization *in vivo* without selecting for resistance

To assess the potential of SMiTE to reduce *S. pneumoniae* colonization, we first evaluated whether repeated exposure to sub-inhibitory concentrations of the bacteriocin could select for bypass mutants. *S. pneumoniae* cultures were passaged daily for 10 passages at 0.25x MIC followed by 10 passages at 0.5x MIC, alongside bacteriocin-buffer and untreated controls. After 20 passages, corresponding to c.a. 170 generations, MIC evaluation revealed no change in susceptibility compared to controls exposed to the bacteriocin-buffer or those that remained untreated, indicating that SMiTE does not readily select for resistant bacteria under these conditions.

We next evaluated whether SMiTE could decrease pneumococcal colonization *in vivo*. Mice colonized with *S. pneumoniae* D39 for three days were treated intranasally every 8 hours with SMiTE, while control mice received bacteriocin buffer. After three days of treatment, nasal lavages were collected, and pneumococcal colonization was quantified by selective plating (**Fig. 5A**). SMiTE-treated mice exhibited a 65-fold reduction in pneumococcal viability compared to buffer-treated controls (**Fig. 5B**). Notably, no differences in body weight were observed between groups throughout the experiment (**Supplementary Fig. S9**).

**Fig. 5.**
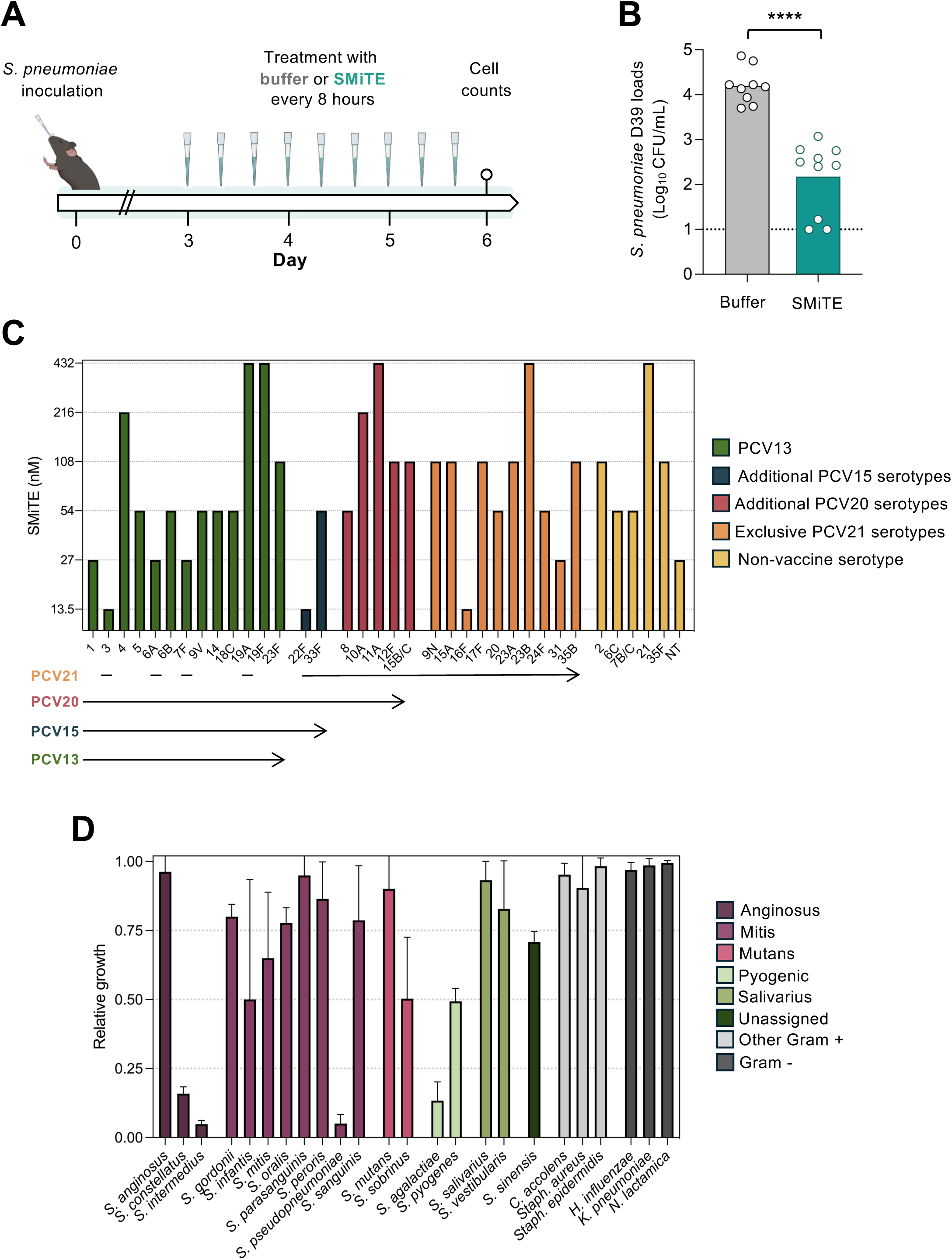
SMiTE reduces pneumococcal colonization *in vivo* and exhibits broad anti-pneumococcal activity with limited effects on other respiratory bacterial species. (A) Experimental design for *in vivo* colonization assay. Mice were colonized intranasally with *S. pneumoniae* D39 for 3 days, then treated every 8 hours with SMiTE or bacteriocin buffer for an additional 3 days (n = 9). Nasal lavages were collected to quantify pneumococcal loads. (B) Quantification of *S. pneumoniae* D39 loads on mice treated with bacteriocin buffer or SMiTE. The two-tailed unpaired Mann-Whitney *U* test was used to compared pneumococcal loads between treatments. **** *P* < 0.0001. (C) MIC evaluation of SMiTE against 36 epidemiologically relevant pneumococcal strains representing both vaccine-targeted and non-vaccine serotypes. Serotypes included in the 13-valent, 15-valent, 20-valent, or 21-valent pneumococcal-conjugate vaccines (PCV13, PCV15, PCV20, and PCV21, respectively) are indicated. (D) Spectrum of activity of SMiTE against 24 non-pneumococcal bacterial species representatives of the oral and upper respiratory tract microbiota, tested at the highest MIC observed for *S. pneumoniae* (432 nM). Relative growth is shown as the ratio of optical density of SMiTE-treated versus buffer-treated cultures. Bars represent the mean, and error bars indicate standard deviation of three independent experiments. Bar colors group species based on either streptococcal classification groups or Gram classification for the other genera.

These results highlight the potential of SMiTE as a novel therapeutic agent capable of reducing pneumococcal colonization *in vivo* while exhibiting a low propensity for resistance development.

### SMiTE exhibits serotype-independent activity against *S. pneumoniae* with limited off-target effects

To further study SMiTE’s potential as a serotype-independent anti-pneumococcal strategy, we evaluated its activity against a panel of 35 epidemiologically relevant *S. pneumoniae* isolates obtained from human nasopharyngeal samples. This collection encompassed both vaccine and non-vaccine serotypes, providing a representative overview of pneumococcal diversity.

SMiTE exhibited inhibitory activity in the nanomolar range across all isolates tested. The lowest MIC values were observed for serotypes 3, 16F, and 22F, while higher MIC were associated with serotypes 11A, 19A, 19F, 21, and 23B (**Fig. 5C**). Other serotypes displayed mostly MICs clustered between 54 and 108 nM, underscoring the consistent potency of SMiTE across diverse pneumococcal strains.

Because bacteriocins often display narrow activity spectra, often limited to closely related species, we next evaluated whether SMiTE followed this pattern. A diverse panel of 24 bacterial species representative of the oral and upper respiratory tract microbiota was assembled, including both commensal and pathogenic taxa from *Streptococcus* and other Gram-positive and Gram-negative bacteria. Since the highest MIC observed for *S. pneumoniae* was 432 nM, this concentration was used to test the susceptibility of non-pneumococcal species.

Among these, *S. constellatus*, *S. intermedius*, *S. pseudopneumoniae,* and *S. agalactiae* were the most susceptible, while moderate inhibition was observed in *S. infantis, S. sobrinus, S. pyogenes*, and *S. sinensis*. The remaining species were minimally affected, suggesting a narrow activity spectrum (**Fig. 5D**).

Collectively, these results demonstrate that SMiTE exerts serotype-independent activity against *S. pneumoniae* while largely sparing other bacterial species of the respiratory microbiota. This selective activity underscores SMiTE’s potential as a targeted agent to reduce pneumococcal colonization.

## Discussion

This study identifies and characterizes SMiTE, a commensal-derived bacteriocin with potent, serotype-independent activity against *S. pneumoniae*. Our findings demonstrate that SMiTE exerts its anti-pneumococcal effects primarily through compromising pneumococcal membrane integrity, and by triggering a coordinated transcriptional stress response and metabolic reprogramming. Notably, SMiTE significantly reduced pneumococcal colonization *in vivo*, did not select for resistant mutants and displayed limited activity against other members of the human oral and upper respiratory tract microbiota. Collectively, these results position SMiTE as a promising candidate for serotype-independent decolonization of *S. pneumoniae* and as a prototype for developing targeted antimicrobials derived from commensal species.

Membrane disruption emerged as a central mechanism underlying SMiTE-mediated killing. Confocal microscopy and ATP leakage assays revealed a dose-dependent loss of membrane integrity, consistent with pore formation and appearance of an external matrix-like material, observed by SEM. This aligns with the mechanisms described for other class II bacteriocins such as lactococcin G, plantaricin EF, and pediocin PA-1, which disrupt bacterial membranes via amphipathic α-helices that insert into lipid bilayers^26–28^. The predicted helical structure of SMiTE and its ability to permeabilize the pneumococcal envelope suggests a similar mechanism, though direct structural confirmation is still needed. Interestingly, imaging suggested preferential damage at septal regions, reminiscent of nisin and mersacidin, which accumulate at cell division zones which are sites of active peptidoglycan synthesis enriched in lipid II^29,30^. Septal regions in pneumococcus correspond to sites of active peptidoglycan and capsule synthesis, where the cell wall and capsule layers are thinner, potentially rendering these areas more accessible to membrane-active peptides^31,32^. Alternatively, a SMiTE receptor may be enriched at the midcell of target cells. The lack of resistance emergence after repeated sub-inhibitory exposure supports this latter hypothesis, as bacteriocins acting on structurally constrained and essential cell envelope components offer limited mutational escape compared to those dependent on variable surface receptors^33,34^. A similar phenomenon has been reported for mersacidin, which, despite strongly inducing the VraSR and upregulating the *vraDE* ABC transporter in *Staphylococcus aureus*, deletion of *vraE* does not confer increased resistance^35^. Similarly, SMiTE’s inability to select for resistant variants suggests that it may act on an essential and structurally constrained target within the pneumococcal envelope.

The transcriptional response to SMiTE exposure corroborates membrane stress as the primary insult. This is supported by ATP leakage and loss of membrane integrity detected followed SMiTE treatment, which indicate collapse of cellular energy. Transcriptomic profiling revealed a coordinated downregulation of gene sets related to translational and carbohydrate metabolism alongside upregulation of two-component-related gene sets. This pattern mirrors canonical bacterial responses to cationic antimicrobial peptides, which activate two-component systems and repress energy-intensive biosynthetic pathways to maintain membrane potential^36–39^. Functional analysis confirmed that gene deletions in *cysK*, involved in sulfur metabolism, and *yvdE* and SPD_0614, involved in cellular metabolic processes, slightly reduced susceptibility to SMiTE, while deletion in *malPQ,* involved in maltodextrin metabolism, slightly enhanced it. These subtle variations in sensitivity indicate that the metabolic state can transiently modulate tolerance, at sub-MIC concentrations, to membrane-targeting agents rather than confer stable resistance. Together, these findings highlight the tight interplay between cellular metabolism and intrinsic resilience to envelope disruption.

The *in vivo* efficacy of SMiTE demonstrates its translational potential. Intranasal administration significantly reduced pneumococcal colonization demonstrating that this bacteriocin retains activity under physiologic conditions and can effectively limit carriage within the host. Its serotype-independent activity can potentially complement a current limitation of pneumococcal conjugate vaccines, which, despite their effectiveness, are constrained by finite valency and serotype replacement^9,10^. Importantly, SMiTE displayed its lowest MIC against serotype 3, one of the most clinically relevant pneumococcal serotypes. Despite being included in current pneumococcal vaccines, serotype 3 remains a major cause of invasive pneumococcal disease worldwide due to its poor vaccine-mediated protection and persistence in carriage^40,41^. This suggests that commensal-derived bacteriocins could complement existing vaccine strategies, offering a new path toward controlling pneumococcal carriage, and hence the likelihood of disease onset, and interrupting transmission chains within the community. While promising, the scalability and delivery in human populations remain to be determined, and further studies will be needed to evaluate pharmacokinetics and long-term effects in more complex microbial communities.

Equally important is SMiTE’s narrow activity spectrum. Unlike broad-spectrum antibiotics that disrupt the healthy microbiota, SMiTE inhibited only a subset of closely related *Streptococcus* species and spared most other oral and respiratory taxa^7,8^. This specificity likely reflects coevolutionary adaptation between commensals and pathogens occupying the same niche, supporting the concept of precision antimicrobials designed to preserve microbial community stability^42–44^. By maintaining commensal that contributed to colonization resistance and ecological balance, SMiTE could indirectly enhance mucosal resilience, a particularly attractive property for prophylactic applications.

In conclusion, SMiTE exemplifies the untapped antimicrobial potential of commensal-derived bacteriocins and introduces a new framework for pneumococcal control that transcends serotype constraints. By combining selective membrane disruption, low resistance potential, and microbiota preservation, SMiTE represents a promising step toward precision antimicrobials designed to modulate colonization rather than broadly eradicate bacteria. Future mechanistic studies and translational evaluations will be essential to determine whether SMiTE or similar peptides can be harnessed safely and effectively to achieve sustained pneumococcal control and reduce global disease burden.

## Methods

### Bacterial strains and growth conditions

Bacterial strains used in this study are listed in **Supplementary Table S6**. All *Streptococcus* spp. used in this study were grown in Todd-Hewitt broth with 0.5% yeast extract (THY). *Staphylococcus aureus* and *Staphylococcus epidermidis* were grown in tryptic soy broth (TSB), while *Corynebacterium accolens* was grown in TSB supplemented with 0.5% Tween80. *Klebsiella pneumoniae* was grown in lysogeny broth (LB) and *Escherichia coli* was grown in either LB or 2xYeast Extract Tryptone (2xYT), depending on the experimental requirement. *Neisseria lactamica* and *Haemophilus influenzae* were grown in C+Y_YB_ supplemented with 20 µg.mL^-1^ hemin and 20 g.mL^-1^ NAD. For the expression of pBAC::8xHis-BlpO_like2_ in *E. coli*, growth medium was supplemented with 100 µg.mL^-1^ ampicillin and, when appropriate, with 500 µM isopropyl β-D-1-thiogalactopyranoside (IPTG).

### Designing of synthetic bacteriocin-gene fragments for cell-free synthesis

Mature bacteriocin sequences were predicted based on conserved amino acid sequences preceding the double-glycine motif (**Supplementary Table S7**). The corresponding nucleotide sequences were codon-optimized for *E. coli* by TwistBiosciences. Additionally, the ribosome-binding site (RBS) of each bacteriocin was optimized to maximize the translation efficiency in *E. coli* using the RBS Calculator v2.1 from Salis Lab^45^. DNA fragments were designed with minimal T7 promoter (TAATACGACTCACTATAG), RBS, initiation codon (ATG), mature bacteriocin gene codon optimized for *E. coli*, a spacer sequence (GCTTTATCTGAGAATATACTCGAA), and T7 terminator (CCCCTAGCCCGCTCTTATCGGGCGGCTAGGGG). All DNA fragments were synthesized by Twist Bioscience.

### Cell-free synthesis of bacteriocins

To evaluate anti-pneumococcal activity, bacteriocins were synthesized using the PUREfrex®2.0 system (GeneFrontier Corporation, Japan). Cell-free synthesis reactions were prepared according to the manufacturer’s instructions and using 10nM as the final concentration of each bacteriocin DNA fragment. As a control, a reaction was set up using the same volume of water instead of DNA. All reactions were incubated at 37°C for 4 hours.

### Evaluation of anti-pneumococcal activity of bacteriocins on *S. pneumoniae* planktonic growth

Cell-free synthesized bacteriocins were tested directly from cell-free reactions, without purification, against *S. pneumoniae* D39 and P537 ^46,47^. For planktonic growth assays, pneumococcal cultures at OD_600nm_ of 0.5 (c.a. 10^8^ cfu.mL^-1^) were diluted to 10^4^ cfu.mL^-1^ in fresh THY containing 1600 U.mL^-1^ catalase and distributed in 384-well plates (90 µL per well). Cultures were treated with 10 µL of cell-free synthesized bacteriocin. Controls received the same volume of water, THY or cell-free reaction without DNA. Growth was monitored for 24 hours by measuring OD_595nm_ every 30 minutes using a plate reader (Tecan Infinite 200 Pro). Three independent experiments were performed for each bacteriocin.

### Evaluation of anti-pneumococcal activity of bacteriocins on solid agar

The inhibitory activity of cell-free synthesized bacteriocins against *S. pneumoniae* D39 and P537 was assessed on solid agar, following a previously described protocol with minor alterations^48^. Briefly, a mixture containing 750 μl of the pneumococcal culture at OD_600nm_ of 0.5, 7125U of catalase, 8 mL of pre-warm THY, and 21ml of molten Tryptic Soy broth with 1% agar was dispensed into three Petri dishes (∼10 mL/plate). After solidification, 2 µL of each cell-free synthesized bacteriocin was stabbed onto the agar. A cell-free reaction without DNA was used as a negative control. Plates were incubated overnight at 37°C in 5% CO_2_. On the following day, plates were inspected for growth inhibition halos around the stabs – indicating inhibitory activity of bacteriocins. Three independent experiments were done for each bacteriocin.

### Heterologous expression and purification of BlpO_like2_

*E. coli* C43 (DE3) cells were transformed with the plasmid pBAC::8xHis-BlpO_like2_ and cultured in 2xYT supplemented with 100 µg.mL^-1^ ampicillin at 37 °C, 180 rpm. When the culture reached an OD_600nm_ of 0.6, protein expression was induced with 500 µM isopropyl β-D-1-thiogalactopyranoside (IPTG) followed by overnight incubation at 30 °C, 150 rpm. Cells were harvested by centrifugation (12,000 x g, 20 minutes, 4 °C) and resuspended in 50 mM Tris, 150 mM NaCl and 1 mM phenylmethylsulphonyl fluoride (PMSF). Lysis was performed using a French pressure cell (16,000 psi) and the lysate was clarified by centrifugation (12,000 x g, 1 hour, 4 °C).

The resulting pellet was washed, sonicated and solubilized in 50 mM K_3_PO_4_, 500 mM NaCl, 6 M Urea, 1 mM DTT, 0.3% Tween 20, 5% glycerol and 20 mM imidazole, followed by ultracentrifugation.

The recombinant 8xHis-BlpO_like2_ was purified from the supernatant using a HisTrap HP 5 mL IMAC column (Cytiva), eluting with a 500 mM imidazole step gradient. Protein-containing fractions were pooled, and buffer exchange was performed using HiTrap desalting columns (Cytiva) into 50 mM K_3_PO_4_, 500 mM NaCl, 0.01% Tween 20 and 10% glycerol to remove the imidazole. The sample was concentrated and sujected to size exclusion chromatography (SEC) on a Hiload Superdex 30 26/600 GL column (Cytiva). Fractions containing purified 8xHis-BlpO_like2_ were pooled, concentrated, and stored at 4 °C.

### Confocal microscopy

The impact of SMiTE on *S. pneumoniae* D39 was evaluated using the LIVE/DEAD viability staining assay or by membrane-staining combined with confocal microscopy. For both assays, *S. pneumoniae* D39 culture at OD_600nm_ of 0.5 was diluted 1:200 (5×10^5^ cfu.mL^-1^) in a final volume of 2 mL containing 1.75 mL was fresh THY and 250 µL of the bacteriocin treatment (0.5x MIC or MIC), buffer, or THY. Growth was stopped 4 hours post-treatment.

Cells were harvested by centrifugation (9,600x g, 3 min), washed three times with 1x PBS and resuspended in 100 µL of 1x PBS. For LIVE/DEAD staining, the cell suspensions were stained using the LIVE/DEAD^TM^ *Bac*Light^TM^ bacterial viability kit, following the manufacturer’s instructions. For membrane staining, cells were stained with 20 µg.mL^-1^ FM4-64 and 5 µM SYTO9 for 15 min at room temperature, protected from light.

Prior to imaging, samples were mounted on a microscope slide with a thin 1.7% agarose pad and visualized using Zeiss LSM 880 point scanning confocal microscope using the Airyscan detector, a 63x Plan-Apochromat 1.4NA DIC oil immersion objective (Zeiss) and the 488nm and 561 nm laser lines. The microscope was controlled using Zeiss v3.0 software which was also used for processing the Airyscan raw images. Three biological replicates were performed for each treatment. Quantitative image analyses were performed using ImageJ/FIJI v2.14^49^.

### Quantification of ATP efflux

To evaluate the impact of SMiTE on *S. pneumoniae*, ATP levels were quantified on pneumococcal cultures exposed to the bacteriocin. For that, *S. pneumoniae* D39 culture at OD_600nm_ of 0.5 was diluted 1:200 to 5×10^5^ cfu.mL^-1^ in fresh THY containing 1600 U/mL of catalase. One hundred and seventy-five microliters of the bacterial suspension were added to each well of a 96-well plate and treated with 25 μL of SMiTE to reach a final concentration of either 0.5x MIC or MIC. As controls, the same volume of bacteriocin buffer or THY was added. Plates were incubated 16 hours at 37°C in 5% CO_2_ atmosphere. After incubation, wells were resuspended and transferred to 1.5 mL microcentrifuge tubes. Cells were harvested by centrifugation (9,600x g, 3 min) and supernatants were filter-sterilized using 0.22 µm filters. After filtration, extracts were diluted 1:10 in ddH_2_O. All samples were kept on ice throughout the preparation process.

ATP levels were measured on cell pellets and supernatants using the ATP Cell Viability Assay (Millipore) according to the manufacturer’s instructions. Luminescence was recorded using BioTek Neo2 Multimode Reader using a gain of 150 and integration time of 10 s. A standard curve of ATP was performed to confirm a linear detection within the range of 0.2 µM to 2 mM ATP.

### Transmission electron microscopy (TEM) and scanning electron microscopy (SEM)

The cellular defects caused by bacteriocins on *S. pneumoniae* D39 cells were examined using both TEM and SEM. The bacterial cultures were treated as described previously for LIVE/DEAD confocal microscopy.

For TEM, cultures were prepared as previously described^50^. Briefly, cells were harvested by centrifugation (800x g, 10 min) and fixed with 2% paraformaldehyde, 2.5% glutaraldehyde, 0.075% ruthenium red and 1.55% lysine acetate. Samples were incubated for 30 min at 4°C and subsequently washed twice with a 0.075% ruthenium red solution. Cells were then resuspended in a solution containing 2% paraformaldehyde, 2.5% glutaraldehyde and 0.075% ruthenium red, and incubated overnight at 4°C. On the following day, the culture was washed three times with the 0.075% ruthenium red solution. Samples were further processed at the Gulbenkian Institute for Molecular Medicine (Oeiras, Portugal). There, cells were rinsed once with 0.1 M sodium cacodylate buffer and twice with distilled water. Dehydration was carried out on ice using a stepwise ethanol series (10%, 30%, 50%, 10 min each step), followed by overnight at 4°C in 2% uranyl acetate prepared in 70% ethanol. Samples were then infiltrated with epoxy resin in increasing ethanol rations (3:1, 1:1, 1:3) and incubated in 100% resin overnight, followed by two additional exchanges with fresh 100% resin. Ultrathin sections were prepared and examined on a Tecnai G2 Spirit BioTWIN transmission electron microscope (FEI) equipped with an Olympus-SIS Veleta CCD camera.

Midcell deformations were noted in a subset of SMiTE-treated cells. To assess the frequency of these alterations, TEM images of SMiTE-treated and control cells were evaluated independently by two researchers blinded to the treatment. Each evaluator was first shown representative examples of cells with and without midcell deformations and then independently scored all images for the presence of these features. The proportion of cells exhibiting midcell deformations was determined by averaging the counts obtained by both researchers.

For SEM, cells were harvested by centrifugation (800x g, 3 min) and washed three times with 1x PBS. Cells were resuspended in a solution containing 1% paraformaldehyde, 2.5% glutaraldehyde and 0.1 M sodium cacodylate and incubated for 30 min at RT. Samples were washed three times with 0.1 M sodium cacodylate. Then, samples were dehydrated using increasing concentrations of ethanol (50%, 70%, 90% and 100%). Each ethanol concentration was incubated for 10 min at RT before removal. Following dehydration, samples were resuspended in tert-butyl alcohol and incubated for 60 min at RT. Cells were collected by centrifugation (800x g, 3 min) and tert-butyl alcohol was added. Samples were frozen at −20°C until further processing. Prior to imaging, samples were lyophilized under vacuum, sputter-coated with gild using a Cressington 108 sputter coater and examined with a Hitachi SU-8010 scanning electron microscope operating at 1.5 kV. All final sample preparation and imaging were performed at the Unidade Militar Laboratorial de Defesa Biológica e Química (Lisboa, Portugal).

### RNA extraction, RNA-Seq and transcriptomic analysis

To evaluate the impact of SMiTE on gene expression, *S. pneumoniae* D39 cultures were treated with either bacteriocin or bacteriocin buffer as a control. Cultures at an OD_600nm_ of 0.5 were diluted 1:200 in 5 mL to achieve an inoculum of 5×10^5^ cfu.mL^-1^ For the treatment, 4.375 mL of fresh THY was mixed with 625 µL of bacteriocin solution, yielding a final concentration of 54 nM (0.5x MIC); for the control, an equivalent volume of bacteriocin buffer was added. Cultures were incubated for 4 hours post-treatment.

Cells were harvested by centrifugation (10,000x g, 5 min) and washed three times with 1x PBS. Transcription was arrested by incubating the pellet in RNAprotect (Qiagen) for 5 min at RT. After centrifugation, supernatants were removed, and pellets were stored at −80°C. For RNA extraction, pellets were thawed on ice, resuspended in 200 µL of 1x PBS and 200 µL of MagNA Pure lysis buffer (Roche), and incubated at 37°C for 1 h. Total RNA was extracted using the *Quick*-RNA Miniprep Kit (Zymo Research) followed by TurboDNase treatment (37°C, 30 min) to remove genomic DNA. Absence of genomic DNA contamination was verified by PCR targeting *gyrA*. A second clean-up step was performed using the same kit. RNA concentration and integrity were assessed using Nanodrop and the Qubit^TM^ RNA IQ Assay Kit, respectively.

RNA sequencing was performed by Novogene (UK) on the Ilumina NovaSeq X Plus platform using a prokaryotic directional mRNA library with rRNA depletion (paired-end, 150 bp read length, 20 million reads per sample).

RNA-Seq analysis was conducted as described by Pobre and Arraiano^51^. Differentially expressed genes (DEGs) were defined as those with false discovery rate (FDR)<0.05, *p*-value<0.05 and log fold-change (logFC)>1 (upregulated) or logFC<1 (downregulated).

Differential expression results containing gene identifiers, logFC, p-value and FDR were further analyzed using R v4.3.1. DEGs were used for KEGG over-representation analysis with the clusterProfiler (v4.10.1) and DOSE (v3.28.2) packages, using *S. pneumoniae* D39 (“spd”) annotation. For gene set enrichment analysis (GSEA), the complete ranked list of all genes was analyzed with the Broad Institute’s GSEA software v4.4.0^52^. Genes were ordered by logFC, and enrichment was tested against KEGG and Gene Ontology (GO) gene sets from the Molecular Signatures Database (MSigDB v2024.1). Enrichment scores were calculated using 1000 phenotype permutations, and pathways with an FDR *q*-value<0.25 were considered significantly enriched.

The core enriched genes identified by GSEA were subsequently analyzed for functional annotation using the GO resource^53,54^. GO term enrichment was performed to identify significantly associated molecular functions, biological processes, and cellular components.

### RT-qPCR validation

To validate the RNA-Seq results, RNA samples were reverse-transcribed into cDNA using the cDNA Synthesis Kit (Bioline). RT-qPCR was performed with the SensiFAST SYBR Kit (Bioline) on a CFX96 Touch real-time PCR detection system (Bio-Rad) using primers listed on **Supplementary Table S6**. *gyrA* was used for normalization. Each sample was analyzed in two technical replicates. Results were considered valid if Ct values between duplicates differed by ≤0.5. Gene expression fold changes were calculated using the 2^-ΔΔCt^ method^55^.

### Construction of deletion mutants

Deletion mutants of all DEGs identified from RNA-Seq analysis were generated by allelic replacement using an antibiotic resistance marker flanked by *lox66* and *lox71* sites ^14,56^. Briefly, the upstream and downstream flanking regions of the target gene(s) were PCR-amplified from the *S. pneumoniae* D39 genome, and the *lox66-KanR-lox71* fragment was amplified from pKan^14^. All fragments were purified using the Zymoclean Gel DNA Recovery Kit (Zymo Research) and assembled via Gibson Assembly (NEB). The assembled construct was subsequently amplified using nested PCR. All primers used for fragment construction are listed in **Supplementary Table S6**.

For transformation, *S. pneumoniae* D39 cultures were grown in C + Y_YB_ medium at 37°C to an OD_600nm_ of 0.5, then diluted 1:100 in fresh medium and grown to an OD_600nm_ of 0.1. At this stage, cultures were supplemented with 200 ng.mL^-1^ of DNA and 100 ng.mL^-1^ of CSP-1 and incubated for an additional 4 hours. Transformants were selected in TSA plates supplemented with 5% sheep blood and 300 µg.mL^-1^ of kanamycin and confirmed by colony-pick PCR.

### Ethics Statement

This study was ethically reviewed and approved by both the Ethics Committee and the Animal Welfare Body of the Instituto Gulbenkian de Ciência (IGC) (license reference A009/2019), and the Direção Geral de Alimentação e Veterinária (DGAV - license reference 0421/000/000/2020), the Portuguese National Entity that regulates the use of laboratory animals. All experiments followed the Portuguese (Decreto-Lei n° 113/2013) and European (Directive 2010/63/EU) legislations, concerning animal welfare, housing, and husbandry.

### Evaluation of SMiTE potential to control in vivo pneumococcal colonization

The *in vivo* inhibitory activity of SMiTE was tested against *S. pneumoniae* D39-Cam^r^ in a mouse model of pneumococcal nasopharyngeal colonization^57,58^. Briefly, six to eight-week-old C57BL/6J female mice were intranasally inoculated, while under general anesthesia with inhaled isoflurane (Abbot), with 10^5^ CFU in 10 µL of *S. pneumoniae*. Mice were daily monitored for clinical symptoms such as weight loss, lethargy, and ruffled fur. Three-day colonized mice were treated intranasally with 10 µL of SMiTE (using a stock at concentration 6.9 μM that corresponds to 0.69 pmol per treatment) or the bacteriocin buffer. Mice were sacrificed with CO_2_ 72 hours after the first bacteriocin treatment and nasal lavage (NL) collected as previously described. Bacterial quantification was performed by 10-fold serial dilution of each NL sample in tryptic soy agar plates with 5% sheep blood supplemented with 5 µg/mL of gentamicin (GBA) to prevent the growth of contaminants.

### Evaluation of minimum inhibitory concentration (MIC)

The MIC of SMiTE was evaluated against a collection of epidemiologically relevant *S. pneumoniae* strains (**Supplementary Table S6**). Pneumococcal strains at OD_600nm_ of 0.5 were diluted 1:200 to 5×10^5^ cfu.mL^-1^ in fresh THY containing 1600 U.mL^-1^ of catalase. One hundred and seventy-five microliters of the bacterial suspension were added to each well of a 96-well plate and treated with 25 μL of two-fold serially diluted bacteriocin. As controls, the same volume of bacteriocin buffer or THY was added. Plates were incubated 24 hours at 37°C with 5% CO_2_ atmosphere. MIC was determined as the lowest concentration of bacteriocin at which no visible growth, by naked eye, was observed after 24-hour incubation. At least three independent experiments were performed.

### Statistical analysis

All statistical analysis were performed using GraphPad Prism v9.0 and R v4.3.1. A binomial generalized linear model (GLM) with Tukey’s Honestly Significant Difference (HSD) post-hoc test was used to compare the proportion of live cells in the LIVE/DEAD assay. Extracellular ATP levels were compared between conditions using the Kruskal-Wallis test followed by pairwise Wilcoxon post-hoc tests with Benjamini-Hochberg correction. For the blinded quantification of septal damage on TEM images, a binomial generalized linear mixed-effects model (GLMM) indicating *Condition* as a fixed effect and *Observer* as a random effect was applied. RT-qPCR validation of RNA-Seq data was analyzed using a ratio paired *t-*test. For *in vivo* colonization data, bacterial loads were compared using the two-tailed unpaired Mann-Whitney *U* test. Differences were considered statistically significant if *p*-value ≤ 0.05.

## Data availability

RNA-seq raw data have been deposited in the ENA database with accession number PRJEB107276.

## Author contributions

R. Sá-Leão and J. Lança contributed to the concept and design of the study. R. Sá-Leão contributed with reagents and materials. Data acquisition was performed by J. Lança, J. Bryton, J. Borralho, C. Candeias and W. Antunes. Data was interpreted by J. Lança, J. Bryton, J. Borralho, C. Candeias, W. Antunes, A. Pandi and R. Sá-Leão. The manuscript was drafted by J. Lança and R. Sá-Leão. All authors critically revised and approved the final version of the manuscript.

## Acknowledgements

We are grateful to Pedro M. Pereira (ITQB NOVA), Sarela Garcia-Santamarina (Hans Knöll Institut), Luisa N. Hiller (Carnegie Mellon University) and Adriano O. Henriques (ITQB NOVA) for fruitful discussions, and the lab of Tobias J. Erb (Max-Planck Institute, Marburg) for hosting J. Lança and J. Borralho during a short visit.

## References

1 Wahl, B. et al. Burden of *Streptococcus pneumoniae* and *Haemophilus influenzae* type b disease in children in the era of conjugate vaccines: global, regional, and national estimates for 2000-15. Lancet Glob Health 6, e744–e757 (2018). 10.1016/S2214-109X(18)30247-X

2 Drijkoningen, J. J. & Rohde, G. G. Pneumococcal infection in adults: burden of disease. Clin Microbiol Infect 20 **Suppl 5**, 45–51 (2014). 10.1111/1469-0691.12461 S1198-743X(14)60175-0 [pii]

3 Collaborators, G. A. R. Global mortality associated with 33 bacterial pathogens in 2019: a systematic analysis for the Global Burden of Disease Study 2019. Lancet 400, 2221–2248 (2022). 10.1016/S0140-6736(22)02185-7

4 Okeke, I. N. et al. The scope of the antimicrobial resistance challenge. Lancet 403, 2426–2438 (2024). 10.1016/S0140-6736(24)00876-6

5 Weiser, J. N., Ferreira, D. M. & Paton, J. C. *Streptococcus pneumoniae*: transmission, colonization and invasion. Nat Rev Microbiol 16, 355–367 (2018). 10.1038/s41579-018-0001-8

6 Simell, B. et al. The fundamental link between pneumococcal carriage and disease. Expert Rev Vaccines 11, 841–855 (2012). 10.1586/erv.12.53

7 Cherazard, R. et al. Antimicrobial resistant *Streptococcus pneumoniae*: prevalence, mechanisms, and clinical implications. Am J Ther 24, e361–e369 (2017). 10.1097/MJT.0000000000000551

8 Langdon, A., Crook, N. & Dantas, G. The effects of antibiotics on the microbiome throughout development and alternative approaches for therapeutic modulation. Genome Med 8, 39 (2016). 10.1186/s13073-016-0294-z

9 Weinberger, D. M., Malley, R. & Lipsitch, M. Serotype replacement in disease after pneumococcal vaccination. Lancet 378, 1962–1973 (2011). S0140-6736(10)62225-8 [pii] 10.1016/S0140-6736(10)62225-8

10 Candeias, C., et al. *Streptococcus pneumoniae* carriage, serotypes, genotypes, and antimicrobial resistance trends among children in Portugal, after introduction of PCV13 in National Immunization Program: A cross-sectional study. Vaccine 42, 126219 (2024). 10.1016/j.vaccine.2024.126219

11 Ganaie, F. A. et al. Update on the evolving landscape of pneumococcal capsule types: new discoveries and way forward. Clin Microbiol Rev 38, e0017524 (2025). 10.1128/cmr.00175-24

12 Narciso, A. R., Dookie, R., Nannapaneni, P., Normark, S. & Henriques-Normark, B. *Streptococcus pneumoniae* epidemiology, pathogenesis and control. Nat Rev Microbiol 23, 256–271 (2025). 10.1038/s41579-024-01116-z

13 Cools, F., Delputte, P. & Cos, P. The search for novel treatment strategies for *Streptococcus pneumoniae* infections. FEMS Microbiol Rev 45 (2021). 10.1093/femsre/fuaa072

14 Borralho, J. et al. Inhibition of pneumococcal growth and biofilm formation by human isolates of *Streptococcus mitis* and *Streptococcus oralis*. Appl Environ Microbiol 91, e0133624 (2025). 10.1128/aem.01336-24

15 Cotter, P. D., Ross, R. P. & Hill, C. Bacteriocins - a viable alternative to antibiotics? Nat Rev Microbiol 11, 95–105 (2013). 10.1038/nrmicro2937

16 Yang, S. C., Lin, C. H., Sung, C. T. & Fang, J. Y. Antibacterial activities of bacteriocins: application in foods and pharmaceuticals. Front Microbiol 5, 241 (2014). 10.3389/fmicb.2014.00241

17 Pandi, A. et al. Cell-free biosynthesis combined with deep learning accelerates de novo-development of antimicrobial peptides. Nat Commun 14, 7197 (2023). 10.1038/s41467-023-42434-9

18 Martin, J. K. et al. A dual-mechanism antibiotic kills Gram-negative bacteria and avoids drug resistance. Cell 181, 1518–1532.e1514 (2020). 10.1016/j.cell.2020.05.005

19 Field, D., Cotter, P. D., Hill, C. & Ross, R. P. Bioengineering lantibiotics for therapeutic success. Front Microbiol 6, 1363 (2015). 10.3389/fmicb.2015.01363

20 Gharsallaoui, A., Oulahal, N., Joly, C. & Degraeve, P. Nisin as a food preservative: part 1: physicochemical properties, antimicrobial activity, and main uses. Crit Rev Food Sci Nutr 56, 1262–1274 (2016). 10.1080/10408398.2013.763765

21 Lopetuso, L. R. et al. Bacteriocins and bacteriophages: therapeutic weapons for gastrointestinal diseases? Int J Mol Sci 20 (2019). 10.3390/ijms20010183

22 Walsh, L., Johnson, C. N., Hill, C. & Ross, R. P. Efficacy of phage- and bacteriocin-based therapies in combatting nosocomial MRSA infections. Front Mol Biosci 8, 654038 (2021). 10.3389/fmolb.2021.654038

23 Huang, F. et al. Bacteriocins: potential for human health. Oxid Med Cell Longev 2021, 5518825 (2021). 10.1155/2021/5518825

24 Mei, M., Estrada, I., Diggle, S. P. & Goldberg, J. B. R-pyocins as targeted antimicrobials against *Pseudomonas aeruginosa*. NPJ Antimicrob Resist 3, 17 (2025). 10.1038/s44259-025-00088-1

25 Sugrue, I., Ross, R. P. & Hill, C. Bacteriocin diversity, function, discovery and application as antimicrobials. Nat Rev Microbiol 22, 556–571 (2024). 10.1038/s41579-024-01045-x

26 Moll, G. et al. Mechanistic properties of the two-component bacteriocin lactococcin G. J Bacteriol 180, 96–99 (1998). 10.1128/JB.180.1.96-99.1998

27 Moll, G. N. et al. Complementary and overlapping selectivity of the two-peptide bacteriocins plantaricin EF and JK. J Bacteriol 181, 4848–4852 (1999). 10.1128/JB.181.16.4848-4852.1999

28 Zhu, L., Zeng, J., Wang, C. & Wang, J. Structural basis of pore formation in the mannose phosphotransferase system by pediocin PA-1. Appl Environ Microbiol 88, e0199221 (2022). 10.1128/AEM.01992-21

29 Brötz, H., Bierbaum, G., Reynolds, P. E. & Sahl, H. G. The lantibiotic mersacidin inhibits peptidoglycan biosynthesis at the level of transglycosylation. Eur J Biochem 246, 193–199 (1997). 10.1111/j.1432-1033.1997.t01-1-00193.x

30 Jensen, C. et al. Nisin damages the septal membrane and triggers DNA condensation in methicillin-resistant *Staphylococcus aureus*. Front Microbiol 11, 1007 (2020). 10.3389/fmicb.2020.01007

31 Nakamoto, R. et al. The divisome but not the elongasome organizes capsule synthesis in *Streptococcus pneumoniae*. Nat Commun 14, 3170 (2023). 10.1038/s41467-023-38904-9

32 Pathak, A. et al. Factor H binding proteins protect division septa on encapsulated *Streptococcus pneumoniae* against complement C3b deposition and amplification. Nat Commun 9, 3398 (2018). 10.1038/s41467-018-05494-w

33 Hasper, H. E. et al. An alternative bactericidal mechanism of action for lantibiotic peptides that target lipid II. Science 313, 1636–1637 (2006). 10.1126/science.1129818

34 Arbulu, S. & Kjos, M. Revisiting the multifaceted roles of bacteriocins: the multifaceted roles of bacteriocins. Microb Ecol 87, 41 (2024). 10.1007/s00248-024-02357-4

35 Sass, P. et al. The lantibiotic mersacidin is a strong inducer of the cell wall stress response of *Staphylococcus aureus*. BMC Microbiol 8, 186 (2008). 10.1186/1471-2180-8-186

36 Majchrzykiewicz, J. A., Kuipers, O. P. & Bijlsma, J. J. Generic and specific adaptive responses of Streptococcus pneumoniae to challenge with three distinct antimicrobial peptides, bacitracin, LL-37, and nisin. Antimicrob Agents Chemother 54, 440–451 (2010). 10.1128/AAC.00769-09

37 Mücke, P. A., Maaß, S., Kohler, T. P., Hammerschmidt, S. & Becher, D. Proteomic Adaptation of *Streptococcus pneumoniae* to the Human Antimicrobial Peptide LL-37. Microorganisms 8 (2020). 10.3390/microorganisms8030413

38 LaRock, C. N. & Nizet, V. Cationic antimicrobial peptide resistance mechanisms of streptococcal pathogens. Biochim Biophys Acta 1848, 3047–3054 (2015). 10.1016/j.bbamem.2015.02.010

39 Assoni, L. et al. Resistance mechanisms to antimicrobial peptides in Gram-positive bacteria. Front Microbiol 11, 593215 (2020). 10.3389/fmicb.2020.593215

40 Horácio, A. N. et al. Serotype 3 remains the leading cause of invasive pneumococcal disease in adults in Portugal (2012-2014) despite continued reductions in other 13-valent conjugate vaccine serotypes. Front Microbiol 7, 1616 (2016). 10.3389/fmicb.2016.01616

41 Luck, J. N., Tettelin, H. & Orihuela, C. J. Sugar-coated killer: serotype 3 pneumococcal disease. Front Cell Infect Microbiol 10, 613287 (2020). 10.3389/fcimb.2020.613287

42 Telhig, S., Ben Said, L., Zirah, S., Fliss, I. & Rebuffat, S. Bacteriocins to thwart bacterial resistance in Gram negative bacteria. Front Microbiol 11, 586433 (2020). 10.3389/fmicb.2020.586433

43 Bomar, L., Brugger, S. D., Yost, B. H., Davies, S. S. & Lemon, K. P. *Corynebacterium accolens* releases antipneumococcal free fatty acids from human nostril and skin surface triacylglycerols. mBio 7, e01725–01715 (2016). 10.1128/mBio.01725-15

44 Brugger, S. D., Bomar, L. & Lemon, K. P. Commensal-pathogen interactions along the human nasal passages. PLoS Pathog 12, e1005633 (2016). 10.1371/journal.ppat.1005633

45 Reis, A. C. & Salis, H. M. An automated model test system for systematic development and improvement of gene expression models. ACS Synth Biol 9, 3145–3156 (2020). 10.1021/acssynbio.0c00394

46 Avery, O. T., Macleod, C. M. & McCarty, M. Studies on the chemical nature of the substance inducing transformation of pneumococcal types: induction of transformation by a desoxyribonucleic acid fraction isolated from pneumococcus type iii. J Exp Med 79, 137–158 (1944).

47 Son, M. R. et al. Conserved mutations in the pneumococcal bacteriocin transporter gene, blpA, result in a complex population consisting of producers and cheaters. mBio 2 (2011). 10.1128/mBio.00179-11

48 Hegde, T. R., Rufus, O. O., Lee, J. & Hong, S. H. Optimizing cell-free protein synthesis for antimicrobial protein production. Methods Mol Biol 2720, 3–16 (2024). 10.1007/978-1-0716-3469-1_1

49. Schindelin, J., et al. Fiji: an open-source platform for biological-image analysis. Nat Methods 9, 676-682 (2012). 10.1038/nmeth.2019

50 Hammerschmidt, S. & Rohde, M. Electron microscopy to study the fine structure of the pneumococcal cell. Methods Mol Biol 1968, 13–33 (2019). 10.1007/978-1-4939-9199-0_2

51 Pobre, V. & Arraiano, C. M. Characterizing the role of exoribonucleases in the control of microbial gene expression: differential RNA-seq. Methods Enzymol 612, 1–24 (2018). 10.1016/bs.mie.2018.08.010

52 Subramanian, A. et al. Gene set enrichment analysis: a knowledge-based approach for interpreting genome-wide expression profiles. Proc Natl Acad Sci U S A 102, 15545–15550 (2005). 10.1073/pnas.0506580102

53 Ashburner, M. et al. Gene ontology: tool for the unification of biology. The Gene Ontology Consortium. Nat Genet 25, 25–29 (2000). 10.1038/75556

54 Aleksander, S. A. et al. The Gene Ontology knowledgebase in 2023. Genetics 224 (2023). 10.1093/genetics/iyad031

55 Livak, K. J. & Schmittgen, T. D. Analysis of relative gene expression data using real-time quantitative PCR and the 2(-Delta Delta C(T)) Method. Methods 25, 402–408 (2001). 10.1006/meth.2001.1262

56 Albert, H., Dale, E. C., Lee, E. & Ow, D. W. Site-specific integration of DNA into wild-type and mutant lox sites placed in the plant genome. Plant J 7, 649–659 (1995). 10.1046/j.1365-313x.1995.7040649.x

57 Valente, C., Cruz, A. R., Henriques, A. O. & Sá-Leão, R. Intra-species interactions in *Streptococcus pneumoniae* biofilms. Frontiers in Cellular and Infection Microbiology 11, 803286 (2022). 10.3389/fcimb.2021.803286

58 Iovino, F., Sender, V. & Henriques-Normark, B. *In vivo* mouse models to study pneumococcal host interaction and invasive pneumococcal disease. Methods Mol Biol 1968, 173–181 (2019). 10.1007/978-1-4939-9199-0_14

59 Janssen, A. B. et al. PneumoBrowse 2: an integrated visual platform for curated genome annotation and multiomics data analysis of *Streptococcus pneumoniae*. Nucleic Acids Res 53, D839–D851 (2025). 10.1093/nar/gkae923

