## Supplementary Figures S1-S9 for "Serotype-independent inhibition of *S. pneumoniae* by SMiTE, a commensal-derived bacteriocin"

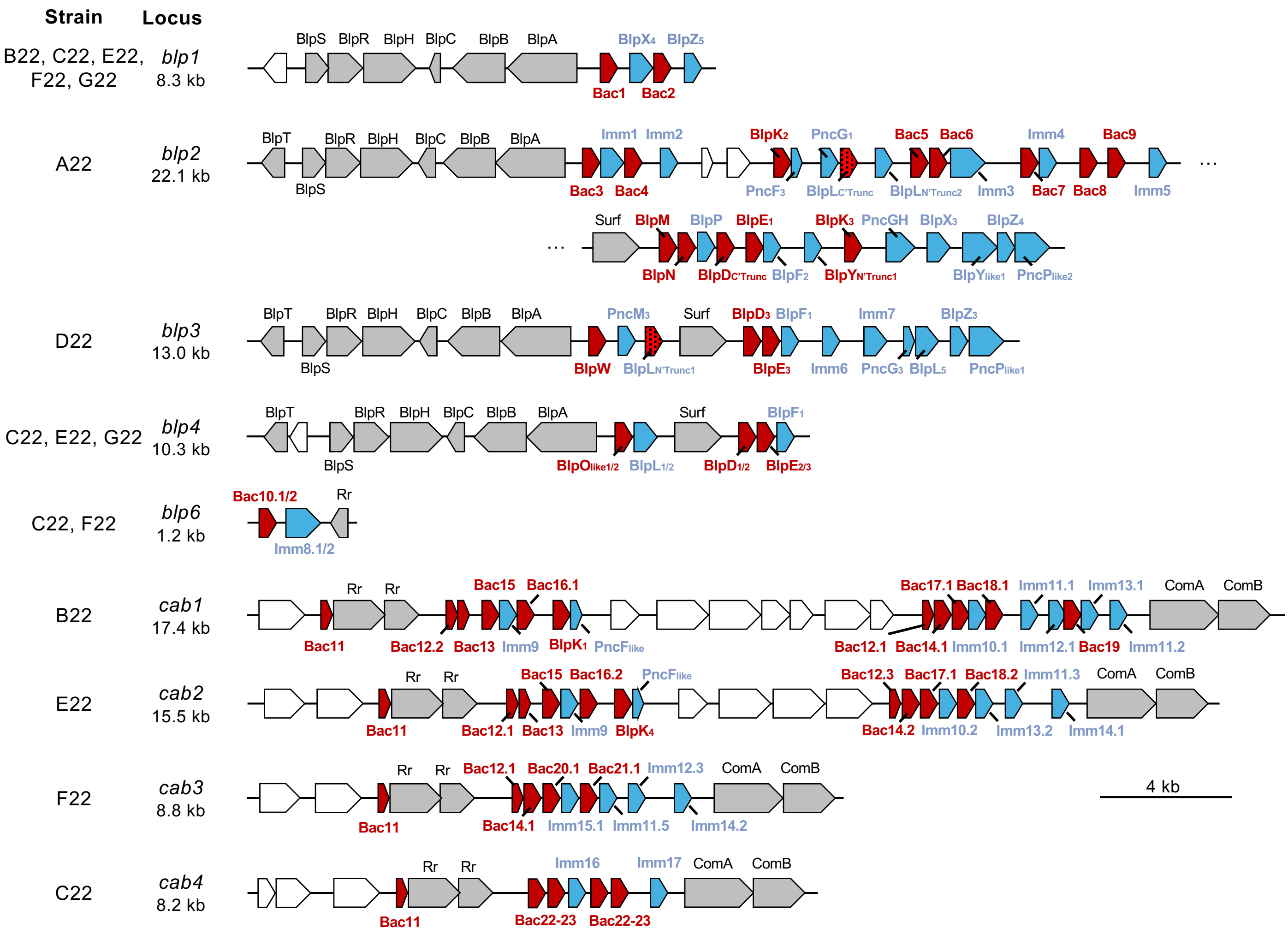

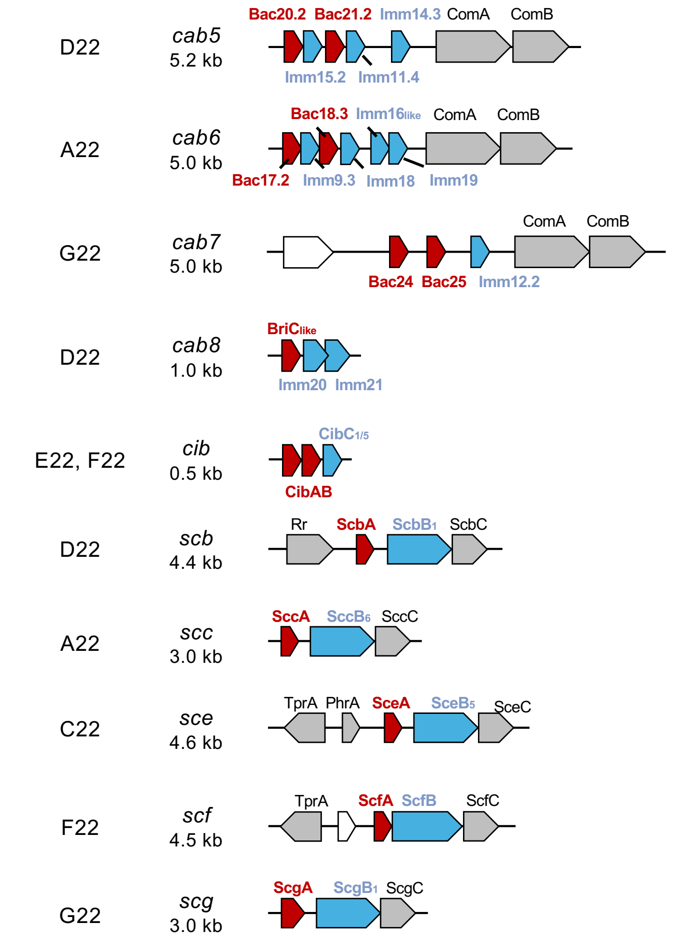


**Figure S1. Genomic context of putative bacteriocin genes and immunity proteins encoded by *S. oralis* strain A22 and *S. mitis* strains B22 to G22.** A schematic representation of all complete bacteriocin-like peptide (*blp*), competence-associated bacteriocin (*cab*) and streptococcin (*sc*) loci from strains A22 to G22 is shown. Putative bacteriocin genes are shown in red. Putative immunity proteins are shown in blue. Bacteriocin accessory genes are shown in grey: BlpT (helicase), BlpS and BlpR (regulator), BlpH (histidine kinase), BlpC (pheromone), BlpA and BlpB (transporters), Rr (response regulator), ComA and ComB (transporters) and ScxC (transporter). Unknown genes and other genes are shown in white. Locus size is indicated next to the locus name. Characterization of these regions has been previously described by us in doi: 10.1128/aem.01336-24.

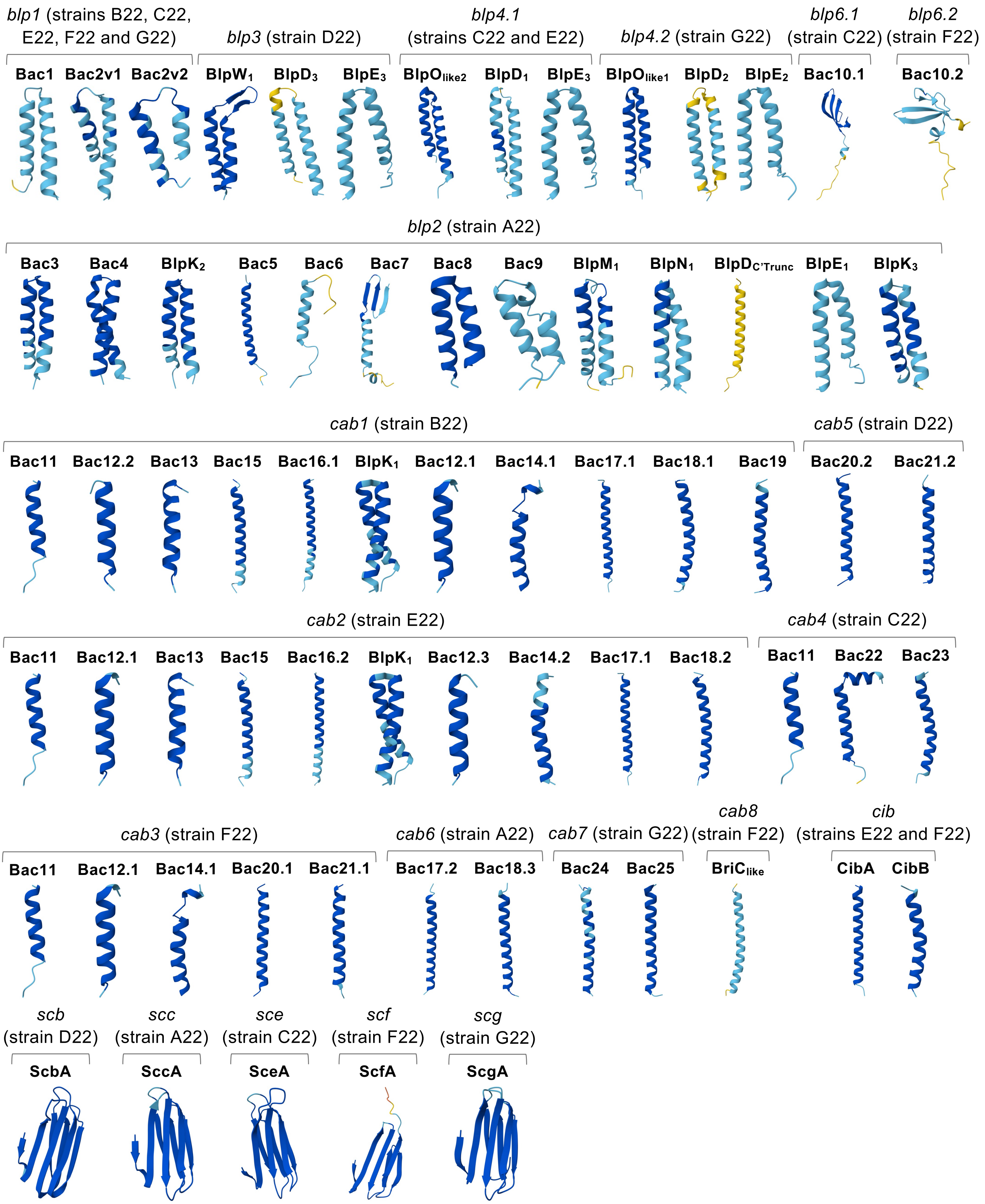


**Figure S2. Predicted structure of bacteriocins from strains A22 to G22 screened for anti-pneumococcal activity.** The monomeric structures of individual bacteriocins were generated using Alphafold (ColabFold v1.5.5) and are colored according to model confidence using the predicted local distance test (pLDDT): orange indicates very low confidence (pLDDT <50), yellow indicates low confidence (50 < pLDDT < 70), light blue indicated confident predictions (70 < pLDDT < 90), and dark blue indicates very high confident predictions (pLDDT > 90). Bacteriocins are grouped according to the bacteriocin loci from which they are encoded.

**A**

**Screening against *S. pneumoniae* D39**

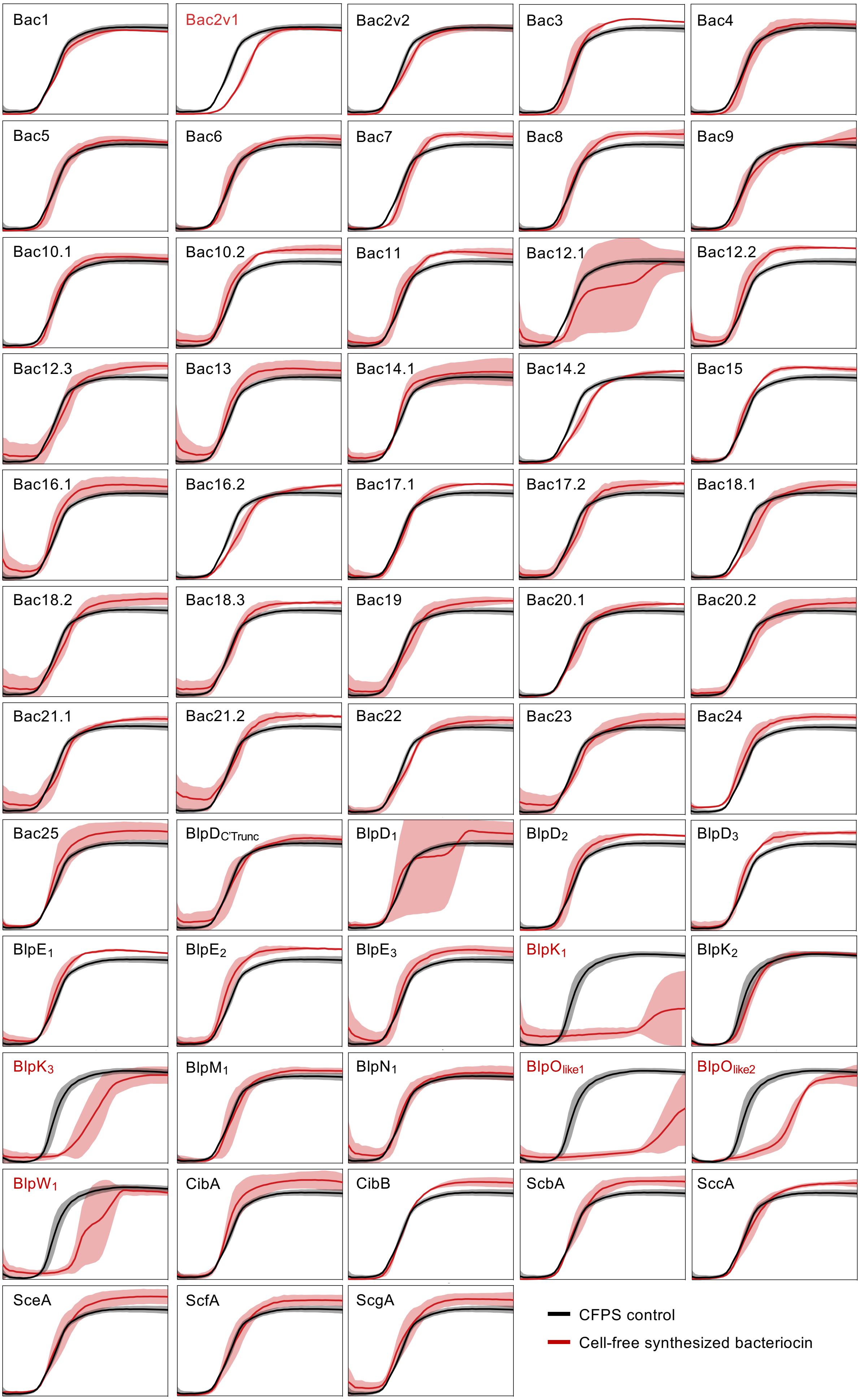




**Figure S3. Screening of 58 bacteriocins from strains A22-G22 against *S. pneumoniae* D39 and P537 using cell-free protein synthesis.**

(continues next page)

**B** **Screening against *S. pneumoniae* P537**

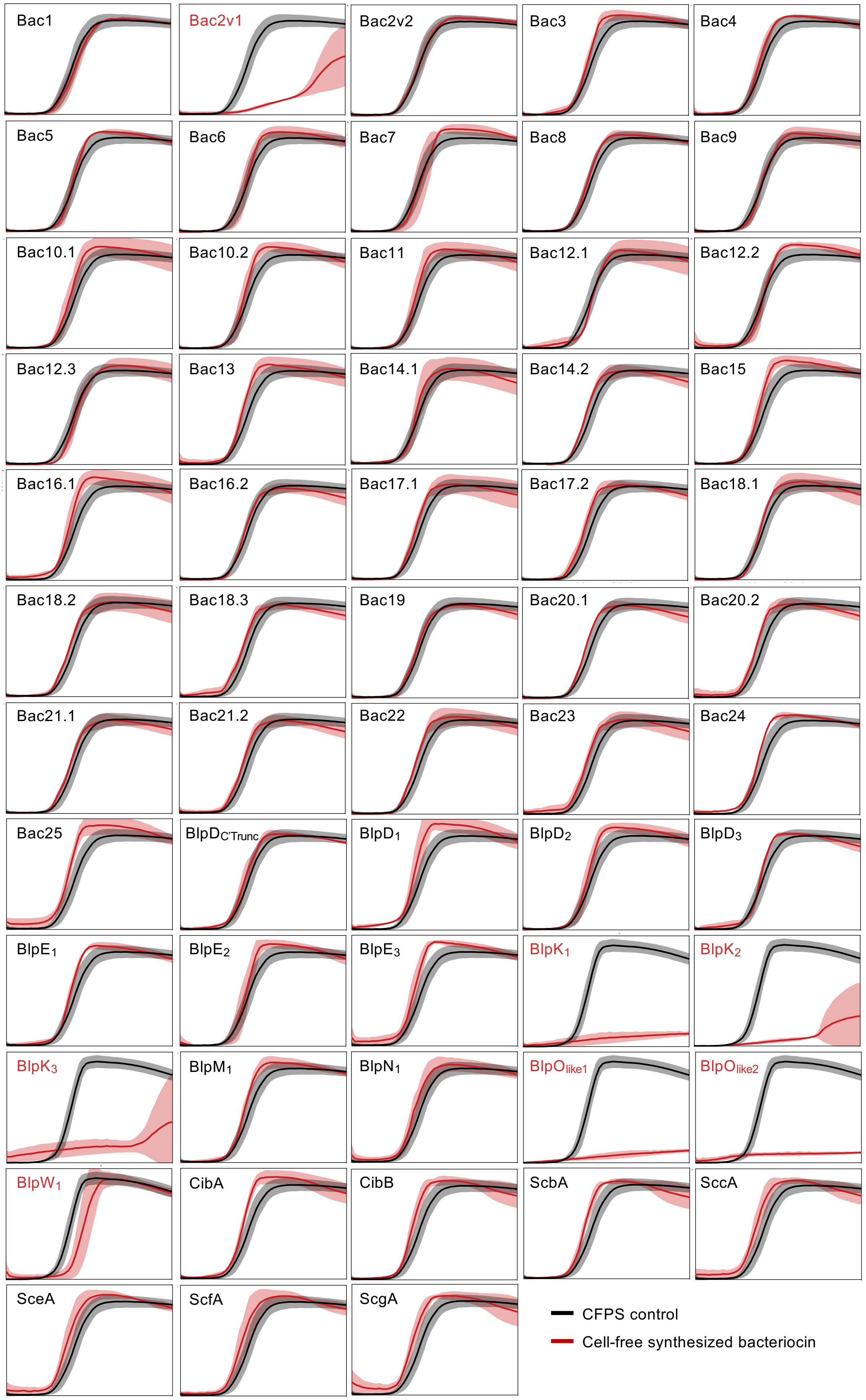




**Figure S3. Screening of 58 bacteriocins from strains A22-G22 against *S. pneumoniae* D39 and P537 using cell-free protein synthesis.**

(continues next page)

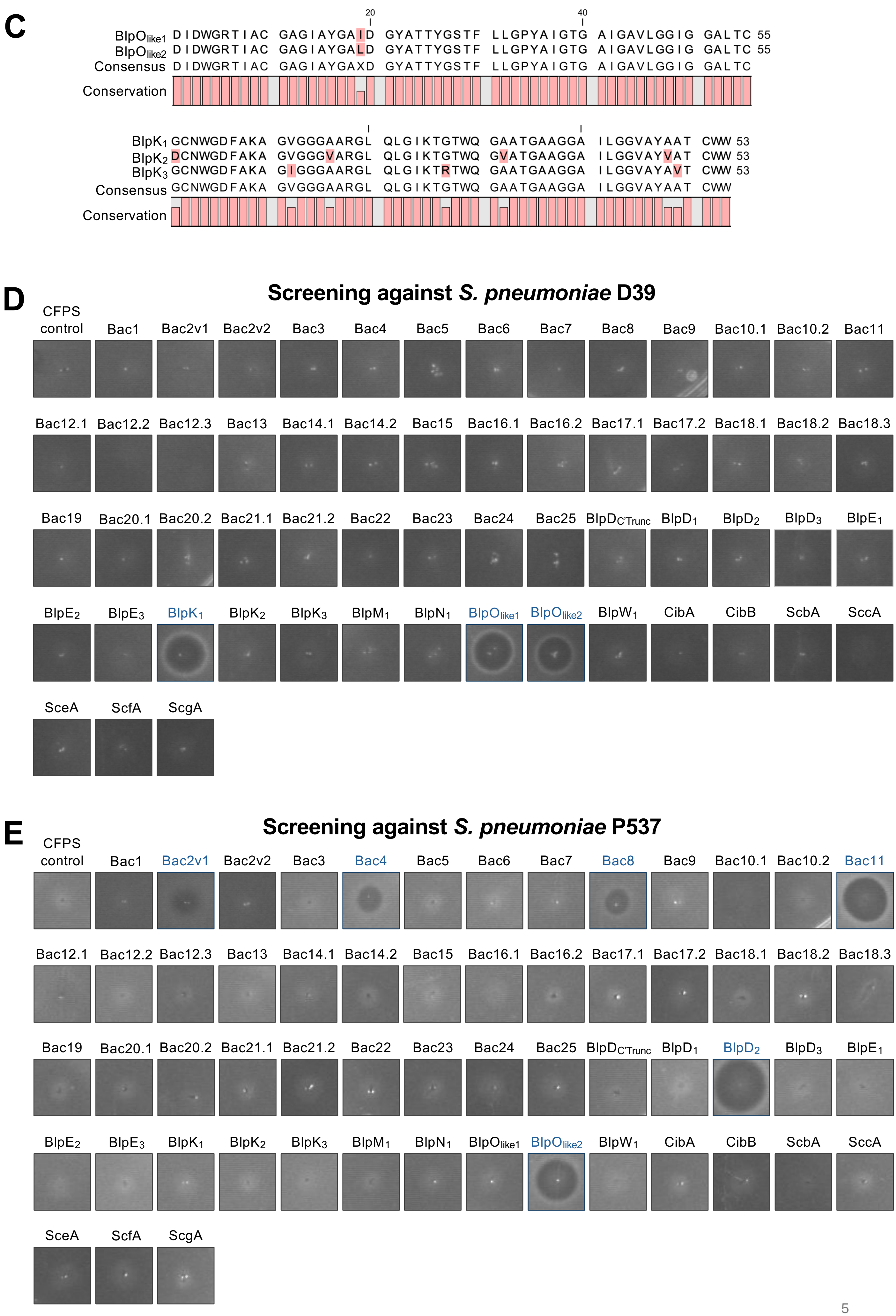




**Figure S3. Screening of 58 bacteriocins from strains A22-G22 against *S. pneumoniae* D39 and P537 using cell-free protein synthesis.**

(legend next page)

**Figure S3. Screening of 58 bacteriocins from strains A22-G22 against *S. pneumoniae* D39 and P537 using cell-free protein synthesis.** (A and B) Growth curves of *S. pneumoniae* D39 (A) and P537 (B) exposed to each of the 58 bacteriocins synthesized using cell-free protein synthesis (CFPS). Black line represents CFPS control, with grey shading indicating the standard deviation. Red line represent bacteriocin-treated cultures, with red shading indicating the standard deviation. (C) Protein sequence alignment of bacteriocins $BlpO_{like1}$ and $BlpO_{like2}$, and of $BlpK_1$, $BlpK_2$ and $BlpK_3$. Alignments were performed using the predicted mature bacteriocin sequences, defined based on double-glycine leader cleavage motifs, and generated with CLC Genomics Workbench 21. (D and E) Representative images of solid media assays showing *S. pneumoniae* D39 (C) and P537 (D) exposed to each of the 58 bacteriocins. Grey outline indicate no inhibition halo, and blue outline indicate the presence of an inhibition halo.

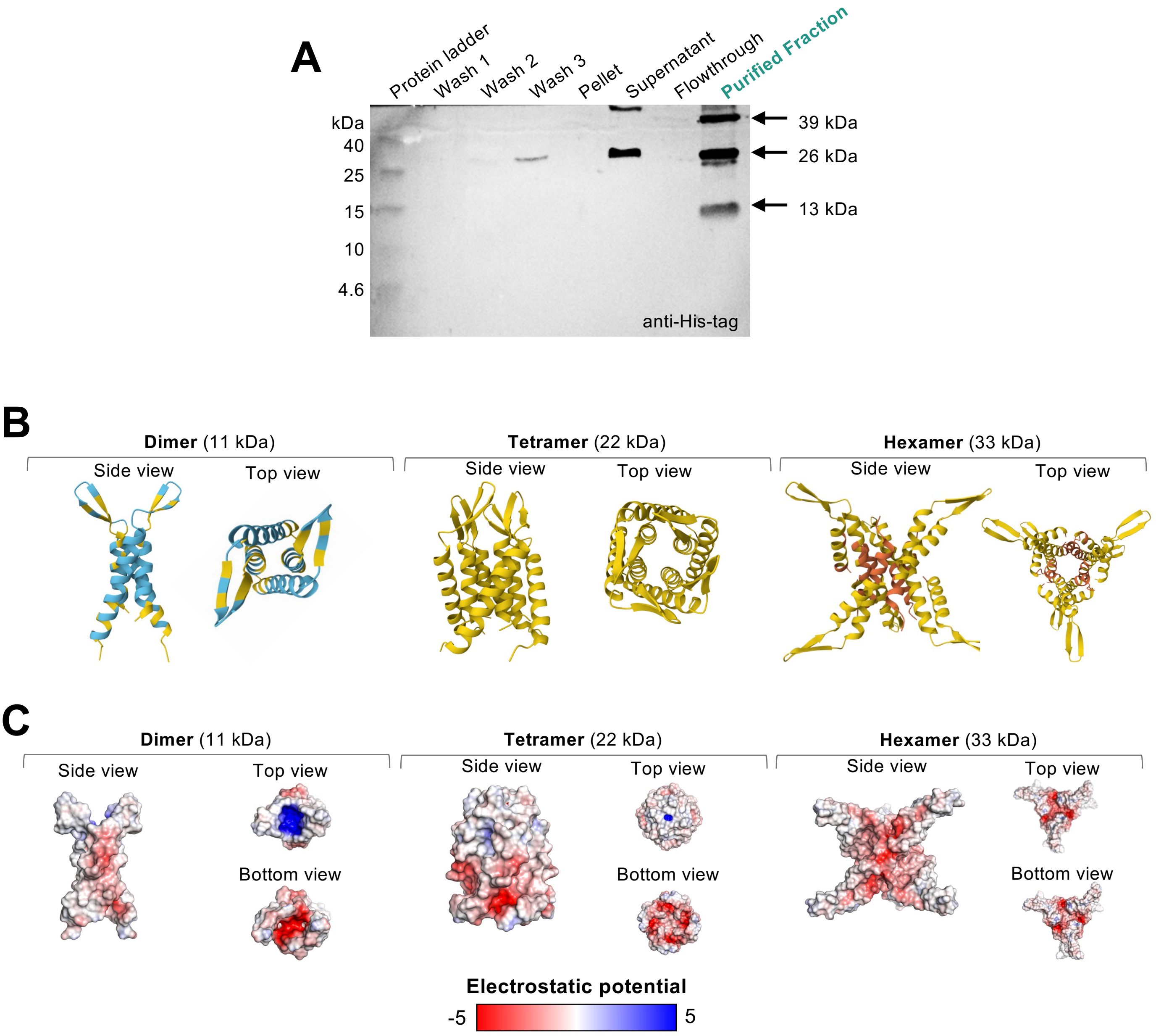


**Figure S4. Purification and structural characterization of native and recombinant SMiTE multimers.** (A) Western blot of samples collected throughout the purification process of SMiTE. Fractions analyzed were wash 1, wash 2, wash 3, pellet, supernatant, flowthrough and purified fraction. Proteins were separated by 16% tricine-SDS-PAGE gel and detected using an anti-His-tag antibody. Three bands were observed at approximately 13, 26, and 39 kDa, which are hypothesized to correspond to dimers, tetramers and hexamers forms of the bacteriocin, respectively. (B) Predicted structure of dimer, tetramer, and hexamer of SMiTE. The structures were generated using Alphafold3 and are colored according to model confidence using the predicted local distance test (pLDDT): orange indicates very low confidence (pLDDT <50), yellow indicates low confidence (50 < pLDDT < 70), light blue indicated confident predictions (70 < pLDDT < 90), and dark blue indicates very high confident predictions (pLDDT > 90). (C) Electrostatic potential structures of dimer, tetramer, and hexamer of SMiTE. Electrostatic potentials were calculated using APBS in PyMOL and mapped onto the molecular surface. Three orientations are presented for each multimer: side, top, and bottom views, to highlight the distribution of charged regions across the assembly. The surfaces are colored on a scale from -5 kT/e (red, negative) to +5 kT/e (blue, positive).

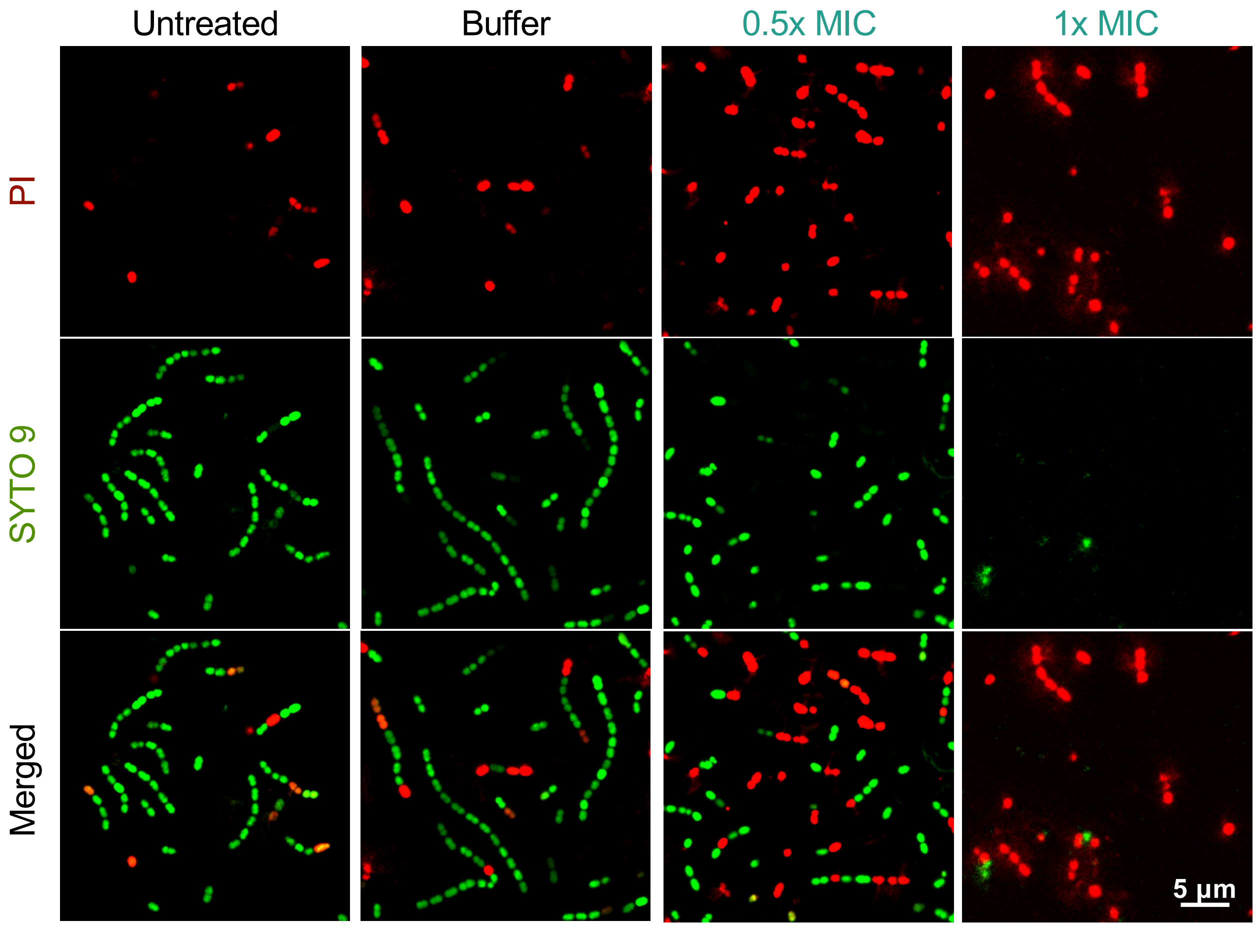


**Figure S5. SMiTE induces dose-dependent membrane permeabilization in *S. pneumoniae*.** Representative confocal fluorescence images of LIVE/DEAD staining of *S. pneumoniae* D39 untreated or exposed to bacteriocin buffer, 0.5x MIC, or 1x MIC of SMiTE. Cells were stained with SYTO9 (intact membranes, green) and propidium iodide (PI) (permeabilized membranes, red). A total of 2689, 2417, 3849, and 1224 cells were analyzed for the untreated, buffer, 0.5x MIC, and 1x MIC conditions, respectively. Images correspond to representative fields from three independent experiments.

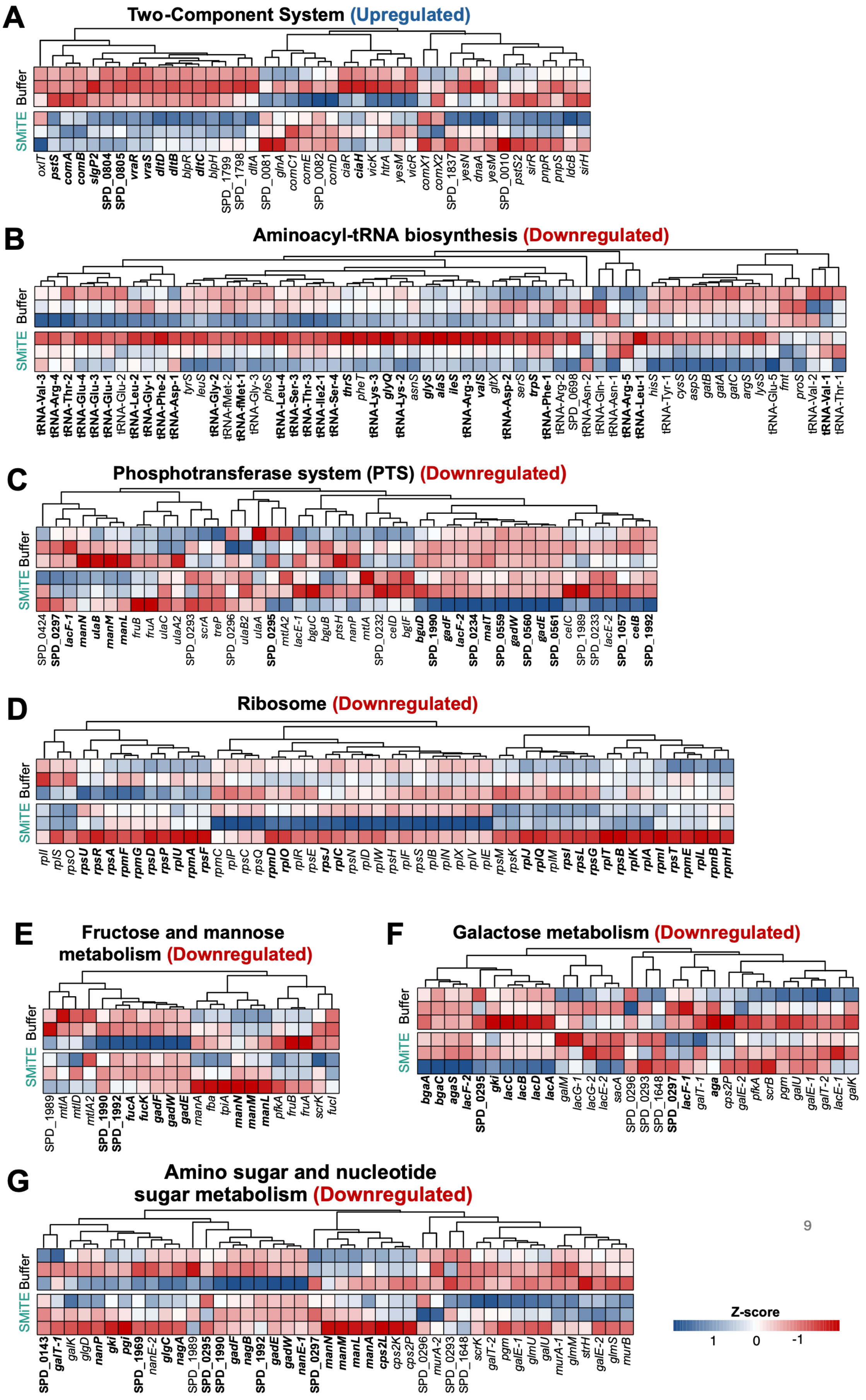


**Figure S6. SMiTE induces coordinated transcriptional remodeling across multiple gene sets and pathways.** Heatmaps showing differential expression (*z*-scores) of genes within gene sets or pathways significantly upregulated (A) or downregulated (B-G) by gene set enrichment analysis (GSEA). (A) Two-component systems; (B) aminoacyl-tRNA biosynthesis; (C) phosphotransferase system; (D) ribosome; (E) fructose and mannose metabolism; (F) galactose metabolism; (G) amino sugar and nucleotide sugar metabolism. Each heatmap shows normalized expression for *S. pneumoniae* D39 cultures treated with 0.5x MIC of SMiTE or bacteriocin buffer (three biological replicates each). Genes in bold correspond to core enriched genes contributing most to the enrichment score. Columns are hierarchically clustered based on similarity of expression patterns. Colors represent relative expression levels (red, lower expression; white, unchanged expression; blue, higher expression).

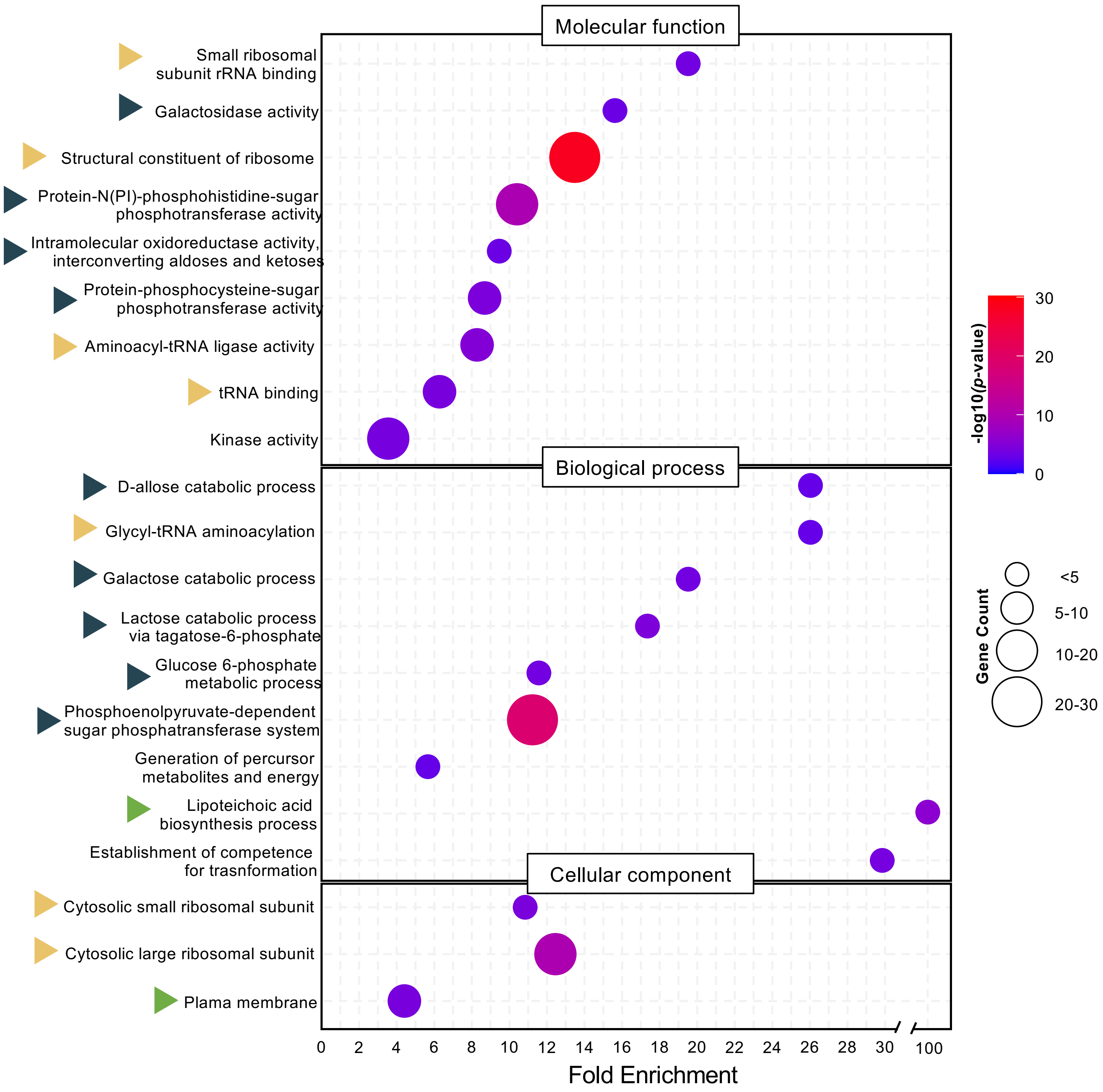


**Figure S7. Gene ontology (GO) analysis revealed that SMiTE exposure enriches functional categories linked to membrane, metabolism, and stress response.** Bubble plots representing GO enrichment of core enriched genes identified by GSEA from *S. pneumoniae* D39 cultures treated with 0.5x MIC of SMiTE compared to bacteriocin buffer control. Enrichment is plotted as a function of fold enrichment. Bubble size corresponds to the number of genes contributing to each GO categories, while color indicates statistical significance ($-\log_{10}$ *p*-value). Blue arrows indicate GO terms related to sugar uptake and carbohydrate metabolism, yellow arrows indicate GO terms related to translation-related processes, and green arrows indicate GO terms associated with membrane and cell envelope remodeling.

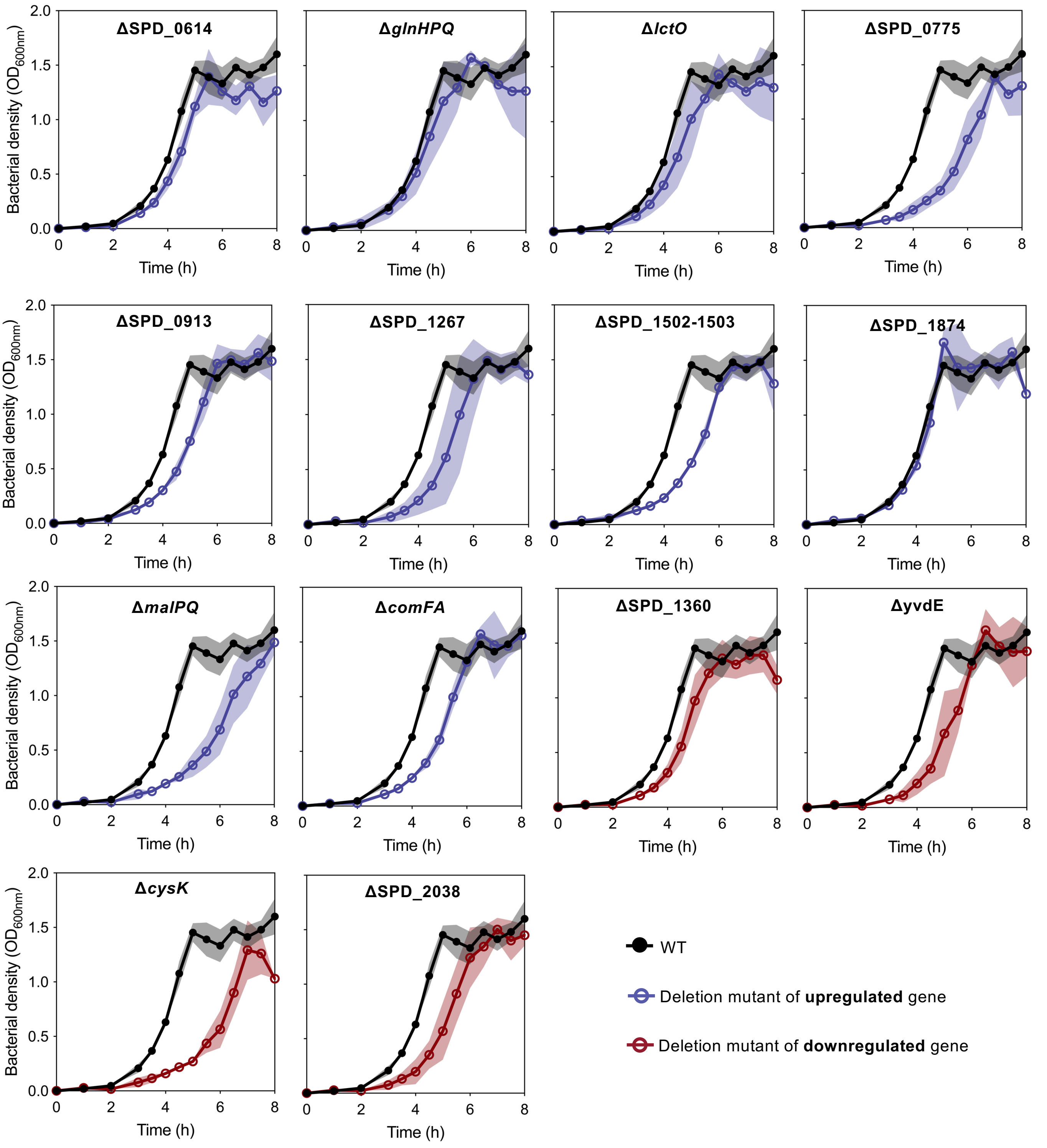


**Figure S8. Growth curves of *S. pneumoniae* of wild-type and isogenic deletion mutants.** Growth curves of wild-type *S. pneumoniae* D39 and deletion mutants of DEGs were monitored in THY. Growth curve of wild-type is shown in black with grey shading for standard deviation; of deletion mutants of upregulated genes are shown in blue with blue shading for standard deviation; and, of deletion mutants of downregulated genes are shown in red with red shading for standard deviation. Cultures were measured every hour until reaching $OD_{600nm}$ of 0.1 and every 30 minutes thereafter until stationary phase.

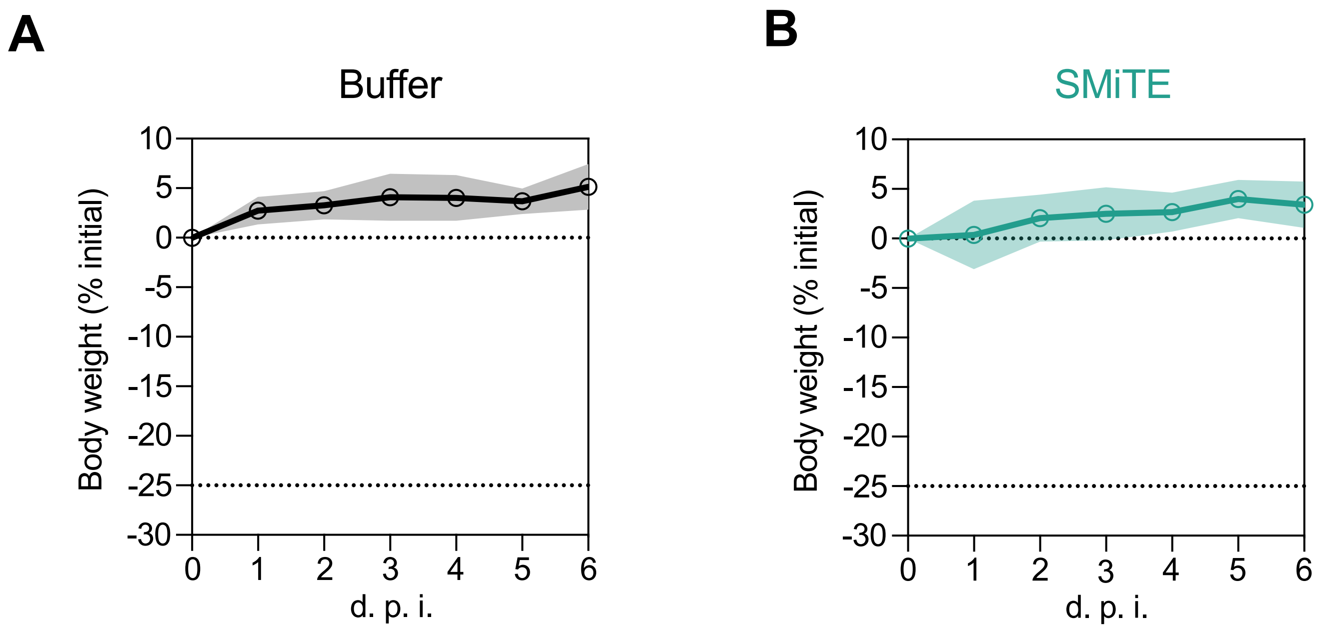


**Figure S9. SMiTE treatment does not affect mice body weight.** Body weight of mice was monitored daily during the *in vivo* colonization experiment and, consequently, during SMiTE treatments. Graphs show the mean percentage change in body weight for each group over the course of the experiment. Standard deviations are represented by shading colors.
